# Siah2 integrates mitogenic and extracellular matrix signals linking neuronal progenitor ciliogenesis with germinal zone occupancy

**DOI:** 10.1101/729152

**Authors:** Taren Ong, Niraj Trivedi, Randall Wakefield, Sharon Frase, David J. Solecki

## Abstract

Evidence is lacking as to how developing neurons integrate mitogenic signals with microenvironment cues to control proliferation and differentiation. We determined that the Siah2 E3 ubiquitin ligase functions in a coincidence detection circuit linking responses to the Shh mitogen and the extracellular matrix to control cerebellar granule neurons (CGN) germinal zone (GZ) occupancy. We found that Shh maintains Siah2 expression in CGN progenitors (GNPs) in a Ras/Mapk-dependent manner. Siah2 supports ciliogenesis in a feed-forward fashion by restraining ciliogenic targets. Efforts to identify GZ sources of Ras/Mapk signaling led us to discover that GNPs respond to laminin, but not vitronectin, in the microenvironment via integrin β1 receptors, which engages the Ras/Mapk cascade, and that this niche interaction is essential for promoting GNP ciliogenesis. As GNPs leave the GZ, differentiation is seamlessly driven by changing extracellular cues that diminish Siah2-activity leading to primary cilia retraction and attenuation of mitogenic responses.

## INTRODUCTION

During neural development, neuronal precursors within germinal zone (GZ) niches simultaneously balance proliferation with the onset of differentiation by responding to morphogens or extracellular matrix (ECM) signaling molecules that have a mitogenic function. A seamless transition of neural precursors from a proliferative state to a terminally differentiated, migratory state underlies the competence of newborn neurons to leave the mitogen-rich GZ to populate the lamina, where these cells will connect with their targets in nascent neuronal circuits(Hatten, 1999). Despite recent progress in understanding the transcriptional cascades and cell-cycle regulatory mechanisms governing the period and output of progenitor proliferation events, relatively little is known about how progenitors process various morphogen and ECM stimuli to integrate responsiveness to mitogens with the onset of GZ exit or subsequent migration.

With its well-defined GZ niche, the cerebellar granule neuron (CGN) is an excellent model with which to study the impact of differentiation transitions on the integration of mitogen responsiveness to GZ exit. Granule neuron progenitors (GNPs) undergo proliferation within the external germinal layer (EGL) and, upon differentiating into CGNs, they migrate radially along the radial glial scaffold, passing through a layer of Purkinje neurons before arriving at their final positions in the internal granule layer (IGL)(Butts et al., 2014). Two cell types provide local and long-range mitogenic signals to GNPs. Pial fibroblasts produce an ECM-rich basement membrane with which GNPs interact via integrin receptors to maintain their proliferation(Blaess et al., 2004; Graus-Porta et al., 2001; Ichikawa-Tomikawa et al., 2012; Pons et al., 2001). Purkinje neurons produce the ligand sonic hedgehog (Shh), a potent mitogen for proliferative GNPs in the outer EGL (oEGL)(Dahmane and Ruiz-i-Altaba, 1999; Lewis et al., 2004; Wechsler-Reya and Scott, 1999). This pathway is frequently deregulated to transform GNPs in a distinct subgroup of cerebellar medulloblastomas (Shh-MBs)(Hatten and Roussel, 2011). The transduction machinery for Shh is localized to the primary cilium, a microtubule-based organelle that is required for the response of GNPs to Shh and, ultimately, for proper cerebellar development(Chizhikov et al., 2009; Satir et al., 2010). Cilium-localized Patched (Ptch) receptor, the negative regulator of the hedgehog pathway, acts as a repressor of Smoothened (Smo), a G-protein–coupled receptor that is the principal activator of the hedgehog pathway(Rohatgi, 2007). Binding of the ligand Shh to Ptch relieves its repression of Smo(Qi et al., 2018), which in turn triggers a series of downstream events that converge into the stabilization and nuclear translocation of the Gli transcription factors(Chen et al., 1999). Importantly, Shh signaling has ceased by the time differentiated CGNs exit the EGL to undergo radial migration, and despite migrating against a gradient of Shh, the CGNs remain insensitive to the ligand. How developing GNPs turn the Shh pathway on or off and how their responsiveness is modified by local cues, such as pia-derived ECM, is unclear and is a key to understanding how normal and transformed GNPs expand during cerebellar development.

CGNs also have a well-characterized cell-intrinsic circuitry controlling GZ exit(Aruga et al., 2002; Famulski et al., 2010; Miyata et al., 1999; Penas et al., 2015; Singh et al., 2016; Trivedi et al., 2017). Prominent among these GZ-exit regulators is the Seven in absentia 2 (Siah2) E3 ubiquitin ligase. This is an evolutionarily conserved cell-fate regulator that is the most downstream component of the Ras GTPase/Map kinase (MAPK) signaling pathway in the specification of R7 cells in the *Drosophila* retina(Carthew and Rubin, 1990). Our laboratory has shown that, during CGN differentiation, Siah2 modulates the acquisition of neuronal polarity and the actin-microtubule interactions necessary for radial migration(Famulski et al., 2010; Trivedi et al., 2017). During the peak stages of progenitor cell expansion, Siah2 expression is high in GNPs, but it is extinguished in CGNs as they exit the EGL. Gain- or loss-of-function studies have shown that Siah2 is necessary and sufficient to maintain GZ occupancy, ultimately by targeting the Pard3 polarity protein and drebrin, the microtubule-actin crosslinking protein, for ubiquitin proteasome degradation. As Pard3 and drebrin act in CGNs to promote radial migration, the relief of Siah2-target inhibition in neuronal differentiation represents a form of integration for cell biological activities linked to GZ exit, positioning Siah2 as a mechanistic entry point through which to explore how GNPs maintain position in their germinal niche. Despite the central role of Siah2 in regulating GNP GZ exit, it is still unclear how extrinsic morphogens and signaling cascades regulating GNP differentiation affect the cell-intrinsic GZ circuity controlled by Siah2 activity or whether Siah2 actively functions to maintain the GNP state.

In this study, we found that, similar to invertebrate Siah orthologues, mouse Siah2 appears to function downstream of Ras/MAPK in CGN GZ exit. We uncovered a surprising link between mitogen signaling and the CGN GZ-exit machinery, as Shh signaling maintains Siah2 expression in proliferative GNPs. We show that the perduring EGL that results from deregulated Shh signaling in GNPs can be rescued by blocking Ras/MAPK and Siah2. We serendipitously discovered that CGNs achieve this effect by retracting their primary cilia and thereby becoming insensitive to Shh, thus enabling them to migrate radially. Interestingly, Ras/MAPK and Siah2 activity maintains primary ciliogenesis in dividing GNPs in part through Siah2 targeted degradation of the Pard3 and Pitchfork proteins. Importantly, we show that the microenvironment surrounding the oEGL, which contains laminin substrates, promotes primary ciliogenesis in proliferative GNPs in an integrin receptor–dependent manner. In our new GZ-exit model, as differentiating GNPs exit their laminin-rich niche, they lose the supportive signals that are required for them to maintain their primary cilia. Siah2 is central to this model because GNPs employ this ubiquitin ligase to maintain GZ occupancy and sensitivity to Shh signaling in a process that is integrated with the engagement of extracellular matrix components in their niche.

## RESULTS

### The Ras-Raf-MAPK pathway controls GZ exit by regulating Siah2

Intrigued by the reported connection between the Ras/MAPK signaling pathway and Siah E3 ubiquitin ligases(Fortini et al., 1992; Schmidt et al., 2007; Simon, 1994), we were curious as to whether this relation was conserved in GNP neurogenesis and GZ exit. We first determined the expression of Ras and phospho-Mek1/2 (pMek1/2) in the developing cerebellum. Immunohistochemical staining for pan Ras and pMek1/2 in combination with Siah2, as a marker for the oEGL, was performed on P7 cerebella from wild-type (WT) mice. Within the EGL, the expression of Ras was uniform throughout, whereas the expression of pMek1/2, a functional readout for active Ras/MAPK signaling, coincided with GNPs having high levels of Siah2 expression. The high level of pMek1/2 staining in the molecular layer seems to originate from the purkinje neurons as judged by their co-localization with Calbindin (**Fig. 1A**). As Siah2 expression in GNPs decreases during postnatal cerebellar development, we wanted to know whether Ras expression followed a similar trend. Ras is a GTPase that cycles between active and inactive states and is capable of signaling to its effectors only when it is in the active state(Bar-Sagi and Hall, 2000). We pulled down the active forms of Ras and performed immunoblot analysis of total Ras and Siah2 in lysates of GNPs isolated from P4, P7, and P10 cerebella, which represent, respectively, the early, peak, and late developmental stages of the cerebellum (**Fig. 1B**). As expected, Siah2 expression decreased from P4 to P10. A similar pattern was seen with total Ras and active Ras. Within the total Ras, the proportion of the active form also decreased from P4 to P10.

**Figure 1.**
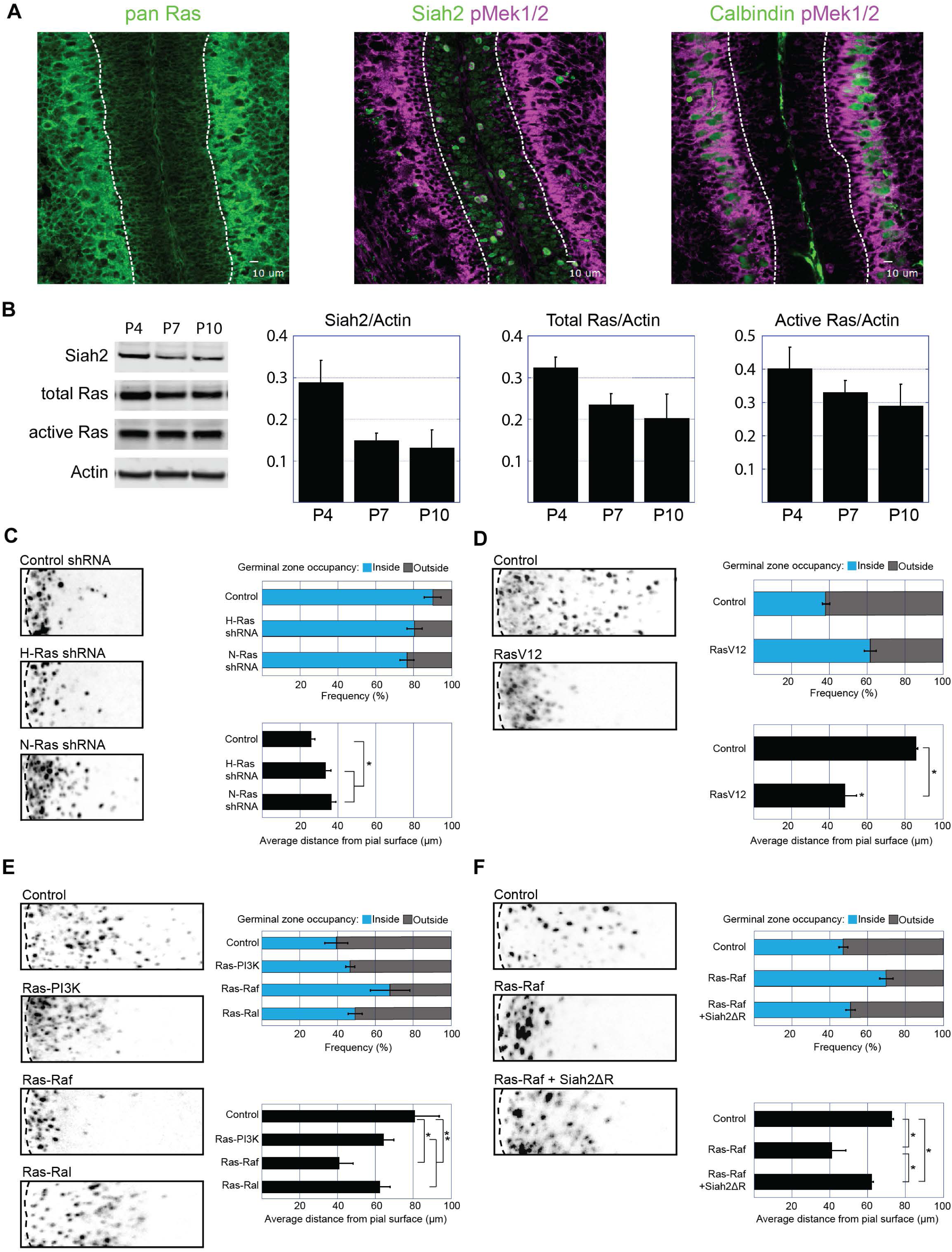
Ras/Mapk controls GZ exit in a Siah2-dependent manner. (**A**) immunohistochemistry staining of pan Ras (left panel), Siah2 and phospho-Mek1/2 co-staining (middle panel), and calbindin and phospho-Mek1/2 co-staining (right panel) in an internal folium of a P7 cerebellum. Dotted lines demarcate the two opposing EGLs. (**B**) Western blot showing the expression of Siah2, total Ras, the active form of Ras, and actin in lysates of isolated GNPs from P4, P7, and P10 cerebella. Western blot quantification is shown in the three graphs at right. (**C–F**) In the following *ex vivo* cerebellar pulse chase assays, GNPs in P7 EGLs were co-electroporated with the indicated constructs and H2B-mcherry. The electroporated cerebella were cultured *ex vivo* for 24 h (C) and 48 h (D–F) before being fixed and imaged. The distance of H2B-labeled cells from the surface of the cerebellum was measured in three independent experiments. The graphs at right show the frequency distribution within and outside the EGL (top panel) and the average migration distance (lower panel). \**P* < 0.05, \*\**P* > 0.05 (by an unpaired two-tailed *t-*test).

Given the evidence that Ras activity diminishes in a temporal window where GNPs mature into CGNs, we wanted to know whether Ras played a role in regulating GZ exit and therefore used a knockdown or gain-of-function approach to study Ras in GNPs. By using the *ex vivo* cerebellar pulse-chase assay developed by our laboratory (**see Materials and Methods**), we can specifically label GNPs and track their migration(Almodovar et al., 2010; Famulski et al., 2010; Singh et al., 2016). Because of the relatively low abundance of K-Ras, we focused our manipulations on H-Ras and N-Ras. The EGLs of P7 WT cerebella were electroporated with miR-30–based shRNAs against H-Ras or N-Ras or against *Renilla* luciferase as a control (**Fig. 1C**). Twenty-four hours post electroporation (hpe), the control cells remained within the EGL, as can be seen by the appearance of a tight band of cells at the surface of the cerebellum. In the H-Ras– or N-Ras–silenced EGLs, however, the cells appeared to begin migrating out of the EGL, with N-Ras silencing having the greatest effect in spurring GZ exit. In contrast, the overexpression of a constitutively active mutant of H-Ras, H-Ras (V12)(Capon et al., 1983), in the EGL (**Fig. 1D**) completely blocked GZ exit at 48 hpe.

Ras triggers the activation of several downstream effector pathways, including the PI3K, Raf-MAPK, and Ral signaling cascades(Cox and Der, 2010). To determine the pathway responsible for controlling GZ exit in GNPs, we electroporated P7 WT cerebella with the constitutively active mutant forms of H-Ras that primarily activate the PI3K, Raf-MAPK, or Ral pathways (**Fig. 1E**). By 48 hpe, most of the control cells had exited the GZ and migrated deep into the cerebellum. The expression of the PI3K- and Ral-activating Ras mutants dampened GZ exit only mildly, whereas the Raf-activating Ras mutant exhibited a greater inhibitory effect that was similar in magnitude to that of Ras V12 (**Fig. 1D**). This suggests that GZ exit controlled by Ras is primarily mediated through the Raf-MAPK pathway. We hypothesized that if Siah2 functioned downstream of the Ras-Raf pathway, then inhibiting its function in the event of constitutive activation of the Ras-Raf pathway should rescue GZ exit. Indeed, by overexpressing a dominant-negative form of Siah2 with a truncated RING domain(Hu and Fearon, 1999) concurrent with the Raf-activating Ras mutant, we could restore GZ exit (**Fig. 1F**). The results of these epistasis experiments suggest that the Ras-MAPK pathway controls GZ exit through Siah2.

### Shh regulates Siah2 in a Ras-Raf-MAPK–dependent manner

Ras-MAPK generally functions with mitogen signaling receptors to regulate cellular proliferation(McCubrey et al., 2007). Having determined that Siah2 functions downstream of the Ras-MAPK pathway in GNP GZ exit, we next wished to identify an appropriate mitogen that interfaced with Ras/MAPK/Siah2 in our system. We focused on the Shh morphogen because the prototypical mitogens that signal through receptor tyrosine kinase receptors to stimulate Ras/MAPK activity (e.g., EGF, IGF, or Fgf) are very poor GNP mitogens, whereas Shh is an order of magnitude more potent as a GNP mitogen than any other morphogen tested to date(Gao et al., 1991; Wechsler-Reya and Scott, 1999). Our decision was bolstered by data from the Pediatric Cancer Genome Project (PCGP), which revealed that in human medulloblastomas (MBs), a group of tumors that arise from GNPs(Gilbertson, 2008), the expression of Siah2 transcripts was highest in the Shh subgroup of the tumors (**Fig. 2A**). To extend the finding of Siah2 expression in a defined genetic context in which Shh signaling is genetically activated in GNPs, we performed immunohistochemical staining for Siah2 on samples of Shh-driven MBs obtained from the Ptc^+/−^ P18^KO^ MB mouse model(Uziel et al., 2005) (**Fig. 2B**). The staining revealed that Siah2 was expressed within the tumor but not in neighboring untransformed cerebellar tissue. To further determine whether Shh signaling regulated Siah2 in untransformed progenitors of the CGN lineage, we grew isolated GNPs from P7 WT cerebella in culture in LacZ- or Shh-N–conditioned medium (**Fig. 2C**). The expression of Siah2 remained unchanged after 24 h. However, at 48 h, the control cells cultured in LacZ-conditioned medium, but not those cultured in Shh-N–conditioned medium, had downregulated Siah2, suggesting that Shh signaling could maintain Siah2 expression. We confirmed this finding by overexpressing the following proteins that activate the Shh pathway in GNPs: Smoothened-M2 (SmoM2)(Xie et al., 1998), a constitutive active mutant of Smoothened; Gli1(Yoon et al., 1998), the transcription factor upregulated in response to Shh pathway activation; and Gli2ΔN(Roessler et al., 2005), a truncated form of Gli2 that acts as a transcription activator; along with LacZ as a control (**Fig. 2D**). All but the LacZ-encoding construct could drive Siah2 expression. We tested whether Shh signaling was required for Siah2 expression by using a cilium-deletion approach in which we silenced Intraflagellar transport protein 88 (Ift88). Shh signal transduction obligately requires a primary cilium, and its depletion via Ift88 or Kif3a loss of function renders GNPs insensitive to Shh signaling(Huangfu and Anderson, 2005; Huangfu et al., 2003; Pazour et al., 2000). Control cells showed a strong increase in Siah2 expression in response to Shh-N stimulation, but this increase was dampened in Ift88-knockdown cells (**Fig. 2E**). We also confirmed that Ift88 knockdown decreased the number of primary ciliated cells in culture (**fig. S2**. Finally, application of small molecule inhibitors of Gli transcription factors also inhibited Shh-N stimulation of Siah2 expression **(fig. S3)**. Taken together, these results show that Shh signaling is necessary and sufficient to maintain Siah2 expression in GNPs.

**Figure 2.**
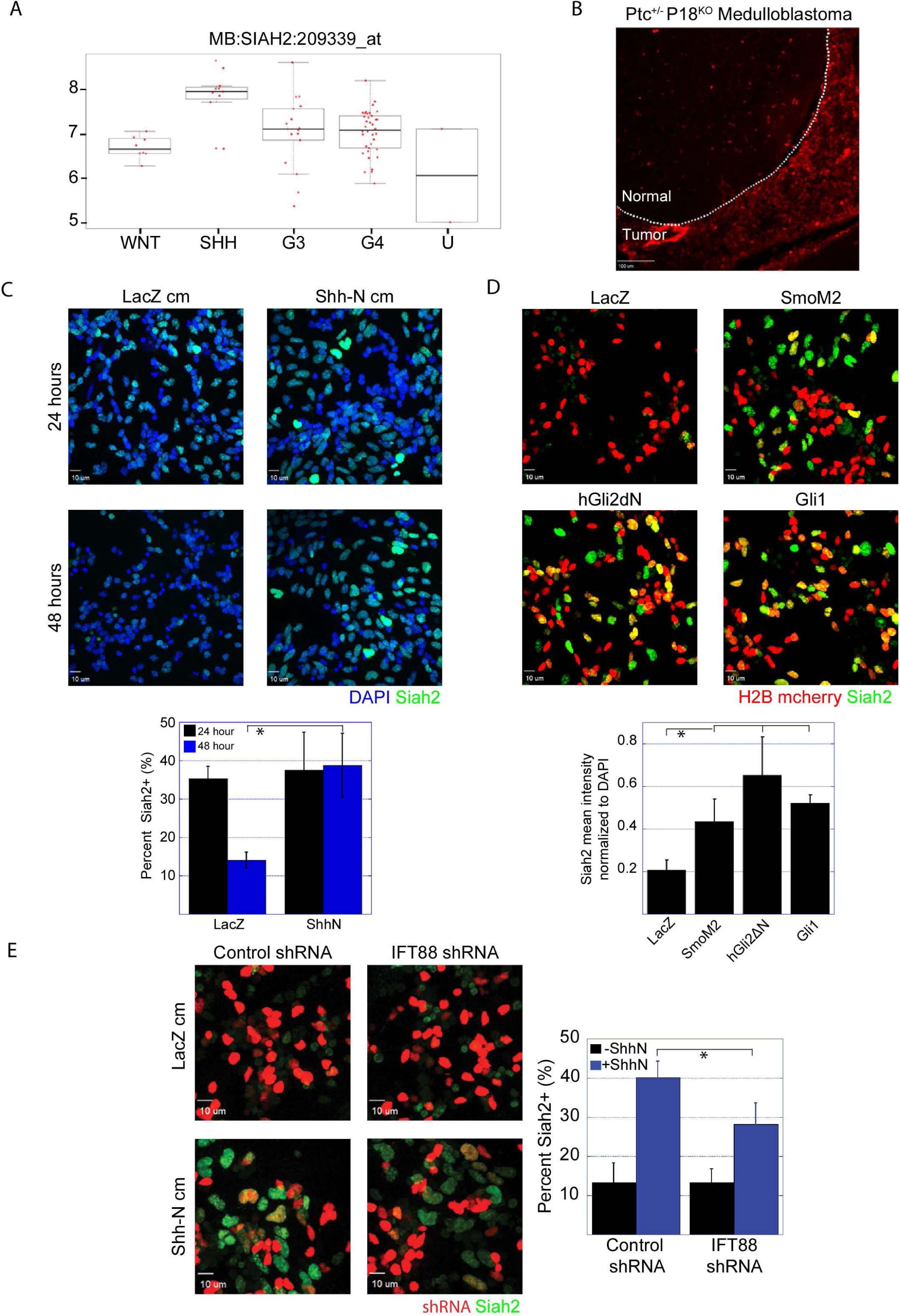
Shh maintains Siah2 expression in a Ras/Mapk-dependent manner. (**A**) Siah2 transcript expression in the different subgroups of human medulloblastomas. U = uncategorized (the subset of tumors that do not belong to any of the four subgroups). Data were obtained from the Pediatric Cancer Genome Project. (**B**) immunohistochemistry staining of Siah2 on a section of mouse Ptc^+/−^ P18^KO^ medulloblastoma tumor. Dotted lines demarcate the boundary between normal and transformed tissue. (**C–E**) In the following experiments, isolated GNPs from P7 cerebella were cultured on Matrigel-coated glass as indicated. cm = conditioned medium. (C) Cells were fixed after 24 and 48 h, and Siah2 immunofluorescence staining was performed. The graph shows the percentage of Siah2-positive cells. (D) Cells were co-nucleofected with the nuclear marker H2B-mCherry and the indicated constructs and cultured for 48 h before being fixed and immunostained for Siah2. The Siah2 mean intensity was measured using SlideBook. Quantification results were as indicated on the graph. (E) Cells were co-nucleofected with the nuclear marker H2B-mCherry and the indicated constructs and cultured for 48 h. They were then fixed, and immunofluorescence staining for Siah2 was performed. The graph shows the percentage of Siah2-positive cells. \**P* < 0.05 (by an unpaired two-tailed *t*-test).

Our epistasis experiments (**Fig. 1**) showed that Siah2 can function downstream of the Ras-MAPK cascade in controlling GZ exit. To determine whether the maintenance of Siah2 expression by Shh signaling required the Ras-MAPK cascade, we first used a pharmacologic inhibitor approach. We stimulated isolated GNPs from P7 WT cerebella with SAG, a small-molecule agonist of the Shh pathway, for 48 h in the presence of vehicle control or one of the following inhibitors of the Ras-MAPK pathway: farnesyl thiosalicylic acid (FTA), a Ras inhibitor; sorafenib (Sor), a Raf inhibitor; trametinib (Tra), a Mek inhibitor; or FR180204 (FR), an Erk inhibitor (**fig. S4**). Pharmacologic inhibition of any component of the Ras-MAPK cascade in the presence of SAG decreased Siah2 expression.

We further tested the dependency of Ras-MAPK signaling on Shh maintenance of Siah2 expression by using our *ex vivo* cerebellar slice system. The EGLs of P7 *Ptch^Flox/Flox^* cerebella were electroporated with plasmids encoding codon-optimized Cre recombinase or its inactive mutant to create cohorts of GNPs with and without strong genetic activation of the Shh signaling cascade; Ptch loss of function relieves its restraint on Smo function, leading to constitutive activation of the Shh pathway. Consistent with the results of our *in vitro* SAG treatment, immunohistochemistry staining for Siah2 revealed that Siah2 expression was increased in *Ptch^Flox/Flox^* GNPs expressing Cre recombinase (**Fig. 3B**). Importantly, Siah2 expression was decreased to control levels by knocking down H- and N-Ras or Erk, showing that Shh maintenance of Siah2 expression in GNPs residing within the EGL niche requires the Ras-MAPK cascade.

**Figure 3.**
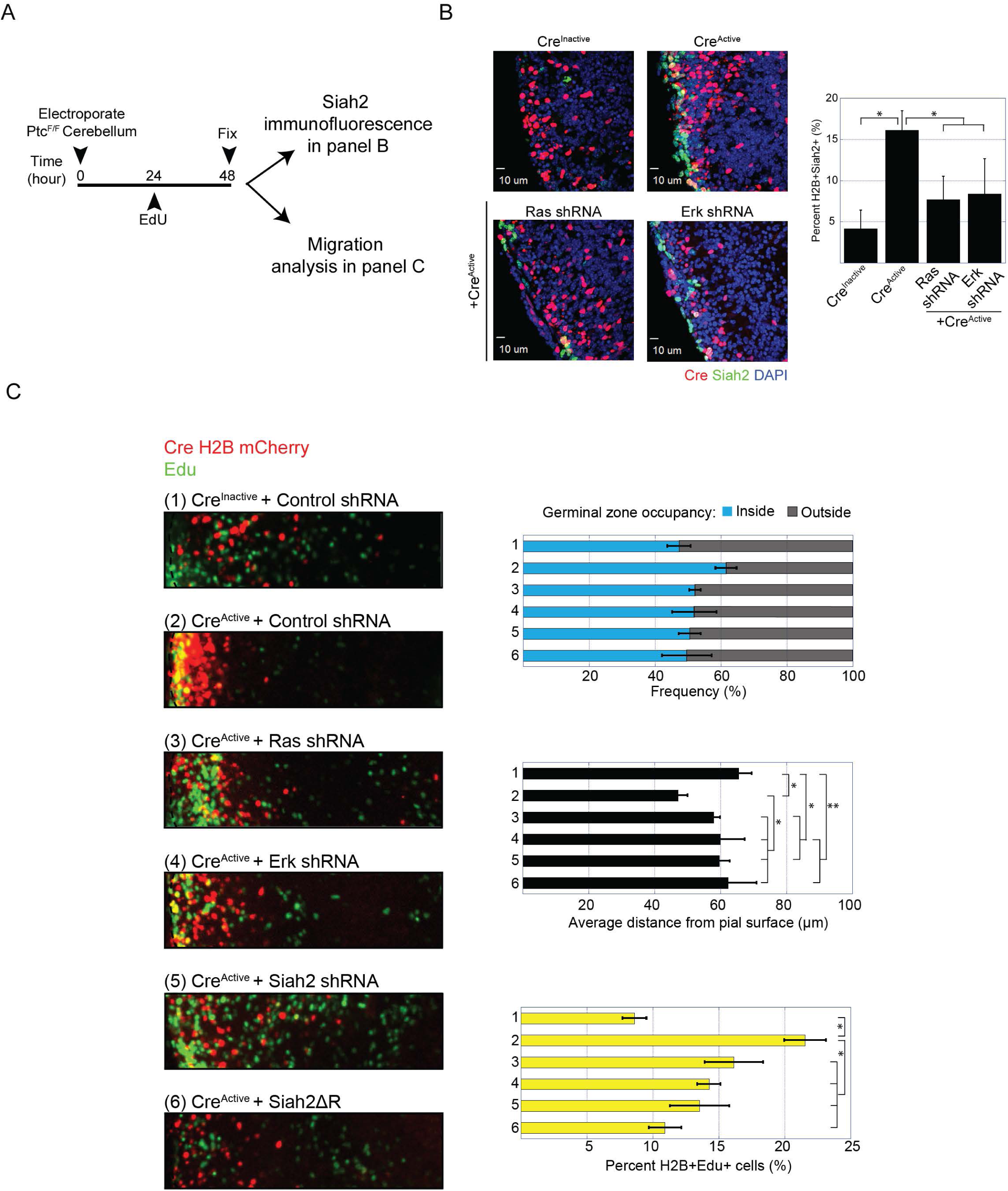
Shh signaling blockds GZ exit in a Ras/Mapk- and Siah2-dependent manner. (**A**) Schematic of the experimental design for panels B and C. GNPs in P7 Ptc floxed (Ptc^F/F^) cerebella were co-electroporated with a bicistronic vector containing Cre recombinase and the nuclear marker H2B mCherry, along with mir30-based shRNA as indicated. Cerebella slices were kept in *ex vivo* culture for 48 h then fixed in 4% PFA in DPBS. (**B**) After fixation, the cerebellar slices were cryoprotected in 30% sucrose and embedded in Neg-50 medium. Sections of 14-μm thickness were prepared and immunohistochemistry staining for Siah2 was performed. Siah2-positive cells among the electroporated cells were quantified, and the results are shown on the graph at right. (**C**) After fixation, the cerebellar slices were stained for EdU by using the Invitrogen Click-it Edu Kit in accordance with the manufacturer’s instructions. Representative images of the migration patterns with the indicated manipulations are shown. The distance of H2B-labeled cells from the surface of the cerebellum was measured in three independent experiments. Electroporated cells that were EdU positive were counted. The graphs at right show the frequency distribution within and outside the EGL (top panel), the average migration distance (middle panel), and the percentage of electroporated cells that were EdU positive (lower panel). \**P* < 0.05, \*\**P* > 0.05 (by an unpaired two-tailed *t-* test).

### Constitutively activated Shh signaling blocks GZ exit, which can be rescued by knocking down Ras, Erk, or Siah2 or by inhibiting the function of Siah2

An overt phenotype of sustained Shh signaling in GNPs is a perduring EGL due to a delay in GZ exit. This has been demonstrated in various Shh-MB mouse models(Goodrich et al., 1997; Oliver et al., 2005; Yang, 2008). Although Shh signaling maintains GNPs in a proliferative and undifferentiated state, the Shh-regulated downstream mechanisms controlling GZ exit remain unclear. Given that Siah2 blocks GZ exit in GNPs by preventing their differentiation and polarization, we postulated that Shh signaling blocked GZ exit by maintaining Siah2 expression in a Ras-MAPK–dependent manner. To test this using the *ex vivo* migration assay, we electroporated EGLs of P7 *Ptch*^Flox/Flox^ cerebella with plasmids encoding codon-optimized Cre recombinase or its inactive mutant and maintained the cerebella slices in *ex vivo* culture for 48 h (**Fig. 3C**). EdU was added in the final 24 h of culture to assess proliferation. GZ exit in GNPs expressing the catalytically inactive mutant Cre recombinase proceeded normally with low incorporation of EdU, demonstrating that cell-cycle exit and differentiation were normal in GNPs with two *Ptch* alleles. In contrast, GZ exit was retarded in GNPs expressing active Cre recombinase, and this was coupled with an increase in EdU incorporation. This finding was expected, because sustained Shh signaling due to the loss of both *Ptch* alleles should maintain GNPs in a proliferative state. The GZ-exit phenotype due to the loss of *Ptch1* can be partially rescued by knocking down H- and N-Ras or by co-expressing the dominant-negative form of Siah2, Siah2ΔR, and it can be fully rescued by knocking down Erk1 and Erk2 or Siah2. Proliferation was decreased under all rescue conditions. These data are consistent with the GZ-exit phenotypes and show that sustained Shh signaling blocks GZ exit in GNPs by maintaining Siah2 expression in a Ras-MAPK– dependent manner.

### Differentiated CGNs retract their primary cilia during GNP differentiation

One potential interpretation of the requirement for Ras/MAPK signaling in Shh maintenance of Siah2 expression was that Ras/MAPK was downstream of the Shh signaling cascade. However, we were unable to detect alterations in Ras/MAPK activity when GNPs were treated with Shh or the Shh-pathway agonist SAG (**fig. S5**). Although we discovered serendipitously that Ras/MAPK/Siah2 signaling constituted a parallel pathway to Shh signaling that regulated GNP ciliation levels (discussed in the next section), the initial foundation of this line of investigation was a comprehensive analysis of ciliation in CGNs and GNPs. We found that GNPs placed in culture in the presence of SAG, the small-molecule agonist of the Shh pathway, maintained their primary cilia after 48 h in culture, whereas non-stimulated controls did not do so (**fig. S6**). This suggests that proliferative GNPs maintain their primary cilia, whereas differentiated CGNs retract their primary cilia.

To further investigate our hypothesis, we used two approaches to examine the primary cilia in the developing cerebellum *in vivo*. First, we performed immunohistochemistry staining on frozen P7 cerebellar sections from WT mice to detect the primary cilium markers ADP ribosylation factor–like GTPase 13B (Arl13b)(Caspary et al., 2007) and adenylate cyclase 3 (Ac3)(Bishop et al., 2007), along with co-staining for Tag1, which marks newly differentiated CGNs(Pickford et al., 1989) (**Fig. 4A**). We constructed a 3D rendering of the EGL based on the confocal images and performed a volumetric analysis of the primary cilia. Staining for both primary cilia markers showed that the primary cilia were more abundant in the oEGL than in the iEGL. To determine the changes in the axonemal volume of the primary cilia within the different layers of the EGL, we examined the EGLs of P7 cerebella by using 3D electron microscopy. The typical primary cilium of a GNP and its 3D-rendered image are shown in **Figures 4B and 4B′**, respectively, and the primary cilium of a CGN and its 3D-rendered image are shown in **Figures 4C and 4C′**, respectively. We manually segmented the primary cilia so as to encompass the basal body to the tip of the axoneme. An example of a 3D-rendered data set of a sampled region of the EGL is shown in **Figure 4D**. Manual segmentation and volumetric analyses of the primary cilia, performed using Amira, revealed that the primary cilia of cells within the oEGL are larger than those of cells in the iEGL. These data suggest that differentiated CGNs retract their primary cilia to become insensitive to Shh and thereby downregulate this pathway to initiate GZ exit.

**Figure 4.**
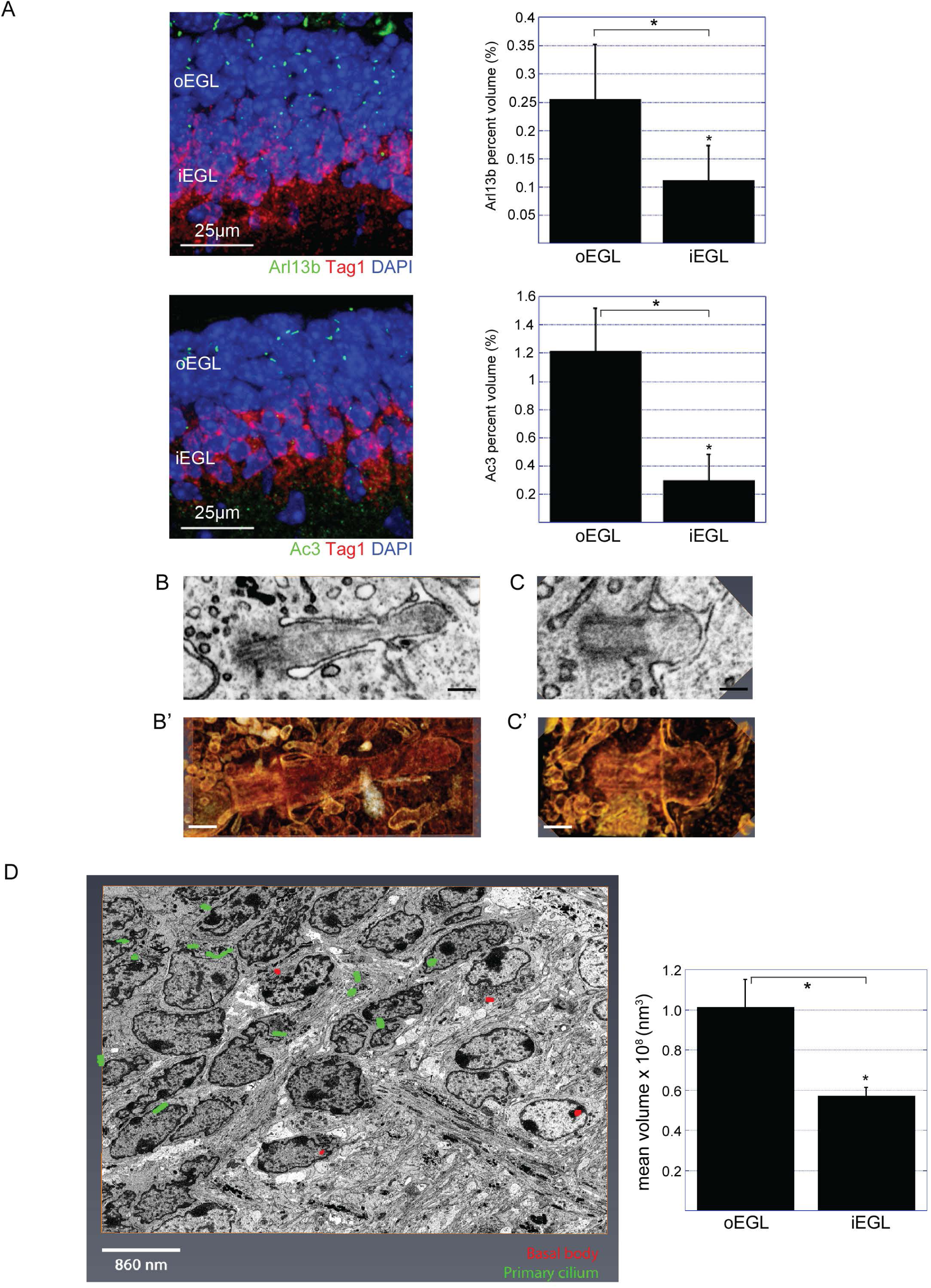
Differentiated CGNs retract their primary cilia during GNP differentiation. (**A**) immunohistochemistry co-staining for the cilium markers Arl13b or Ac3, together with a marker for newly differentiated CGNs, Tag1, in fixed and frozen P7 cerebella sections. Confocal stacks (10 μm) were obtained, and volumetric analyses for Arl13b or Ac3 were performed using Amira. The abundance of cilia within the oEGL and iEGL was represented as the percentage volume within the layers of the EGL, as indicated in the corresponding graphs at right. (**B and B′**) Representative SEM images of a single plane through the primary cilium of a GNP (B) and the corresponding 3D rendering (B′). (**C and C′**) Representative SEM images of a single plane through the primary cilium of a CGN (C) and its corresponding 3D rendering (C′). Scale bars =250nm. (**D**) Representative maximum projected SEM image of the EGL in a P7 cerebellum. The segmentation represents the ciliary axoneme (in green) and basal bodies (red). The graph shows the results of volumetric analysis of the primary cilia within the oEGL and iEGL. \**P* < 0.05, \*\**P* > 0.05 (by an unpaired two-tailed *t-*test).

To determine whether ciliation status had any consequence for GNP GZ occupancy, we electroporated shRNA constructs against Ift88 and *Renilla* luciferase into the EGLs of P7 WT cerebella (**fig. S7**). After 24 h, we detected early GZ exit when we knocked down the primary cilia. The decreased EdU incorporation with Ift88 knockdown demonstrated that the primary cilia were efficiently knocked down, as proliferation is a direct response to Shh signaling in GNPs. Taken together, these data show that GNPs require their primary cilia in order to maintain GZ occupancy.

### Primary ciliogenesis in GNPs in response to Shh requires Ras, Erk, or Siah2, and the loss of primary cilia induces early GZ exit

Given that knockdown of Ras or Siah2 induces early GZ exit, we wanted to know whether the Ras-MAPK pathway or Siah2 had a role in regulating the primary cilia. We nucleofected isolated GNPs from P7 WT cerebella with miR30-based shRNA constructs targeting H-Ras and N-Ras, Erk1 and Erk2, Siah2, or *Renilla* luciferase as a control, and we grew them in culture with or without Shh-N (**fig. S8**). After 48 h, the percentage of ciliated GNPs detected by Arl13b immunocytochemistry was higher in Shh-N–treated cultures than in untreated control cultures. The percentage of ciliated cells was significantly decreased when Ras, Erk, or Siah2 was knocked down, showing that the maintenance of primary cilia in response to Shh requires Ras, Erk, and Siah2.

To determine whether primary ciliogenesis in response to Shh required Ras or Siah2, we developed a confocal time-lapse microscopy assay to visualize this process specifically (**Fig. 5A**). In this assay, we nucleofected GNPs isolated from P7 WT cerebella with constructs encoding venus tagged Arl13b, the FUCCI fragment of human geminin(Sakaue-Sawano et al., 2008), an S-G2-M cell-cycle marker, and nuclear BFP–tagged shRNA. Cells expressing all three constructs were selected at the beginning of the time-lapse study and tracked for 14 h, during which time we could monitor the cells as they underwent cell-cycle progression, mitosis, and primary ciliogenesis. We used this assay to determine mechanistically whether Ras or Siah2 silencing affected primary ciliogenesis. In control GNPs, in the absence of Shh-N, a high proportion of the cells failed to undergo primary ciliogenesis after cell division, whereas ciliogenesis was greatly increased in the presence of Shh-N, showing that Shh signaling promotes GNP primary ciliogenesis. When Ras or Siah2 were silenced in the presence of Shh-N, the proportion of cells that failed to undergo primary ciliogenesis increased relative to that in control Shh-N–treated cells, showing that Shh signaling–driven primary ciliogenesis requires Ras or Siah2.

**Figure 5.**
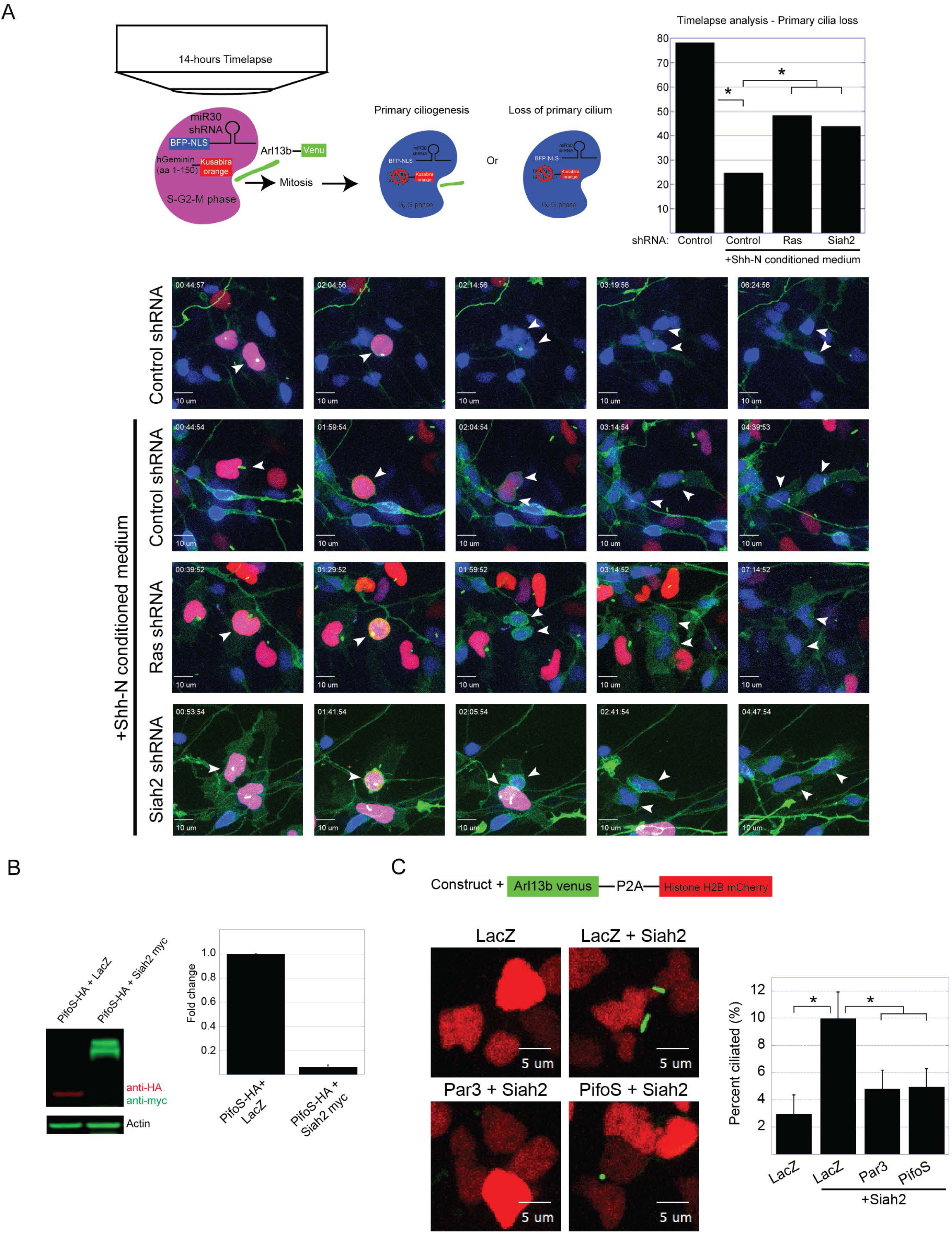
Shh-driven primary ciliogenesis requires Ras and Siah2 inhibition of Pifo and Par3. (**A**) Schema of the experimental design. Isolated P7 CGNs were nucleofected with a combination of the indicated expression vectors and cultured overnight in the presence or absence of Shh-N. Cells expressing all three vectors (BFP-NLS–tagged shRNA, Kusabira orange–tagged geminin, and Venus-tagged Arl13b) were selected, and time-lapse confocal imaging (at 7-min intervals) for 14 h was set up. Cells were scored according to whether (1) either of the progenies, (2) both progenies, or (3) none of the progenies regrew the primary cilium. The graph shows the percentage of cells that did not regrow the primary cilium in each of the indicated conditions. Representative figures from the time-lapse imaging are shown. \**P* < 0.01 (by a χ^2^ test). (**B**) Western blot analysis of Pifo stability. Pifo was co-transfected with LacZ or Siah2 into HEK293T cells, which were then cultured for 24 h. Lysates were then collected and processed for Western blot analysis. The graph shows the results of the quantification of Pifo abundance. (**C**) Isolated P7 CGNs were co-nucleofected with the indicated constructs and the bicistronic vector carrying Venus-tagged Arl13b and mCherry-tagged histone H2B and cultured *in vitro* for 48 h. Live-cell imaging was performed and scored for the proportion of nucleofected cells (H2B-mCherry+) that were ciliated. \**P* < 0.05 (by an unpaired two-tailed *t-*test).

We next focused on understanding how Siah2 controlled ciliogenesis in GNPs. Siah2 specifically recognizes a well-defined degron motif(House et al., 2003) in its targets; thus, we reasoned that Siah2 could potentially promote primary ciliogenesis by targeting proteins that promote cilium disassembly for degradation. Therefore, we scanned the literature for candidate proteins that played a role in cilium disassembly and contained Siah2 degron motifs, with a view to using them in a screen to complement ciliation phenotypes related to the activation of Shh signaling or Siah2 gain of function. We identified four candidates: the F-actin cross-linking protein α-actinin 4 (Actn4); the actin binding protein drebrin (Dbn)(Nager et al., 2017); a microtubule-depolymerizing kinesin (Kif19a)(Niwa et al., 2012); and the protein encoded by the mouse embryonic node gene *Pitchfork* (Pifo), which exists in two isoforms: the full-length isoform (PifoL) and a shorter isoform (PifoS) (Kinzel et al., 2010). We generated constructs encoding each Siah2 target protein and nucleofected GNPs isolated from P7 WT cerebella in conjunction with the bicistronic vector encoding a Venus-tagged Arl13b and mCherry-tagged histone H2B, then we grew the cells in culture in the presence or absence of Shh-N. Also included in the screen was the polarity protein Par3, a known target of Siah2 (**fig. S9**). The bicistronic construct enabled us to perform live-cell imaging and to determine the percentage of ciliated cells among the nucleofected cells. In this screen, we identified PifoS and Par3 as targets of Siah2 that could block Shh-driven primary ciliogenesis. To confirm that PifoS was a target of Siah2, we co-expressed it with Siah2 in HEK-293T fibroblasts (**Fig. 5B**) and found that PifoS was diminished in the presence of Siah2.

Next, we investigated whether Siah2 was sufficient to maintain the primary cilia in GNPs. We nucleofected GNPs isolated from P7 WT cerebella with the bicistronic vector encoding the Venus-tagged Arl13b and mCherry-tagged histone H2B and constructs encoding Siah2 in combination with LacZ, Par3, or PifoS, and we maintained the cells in culture for 48 h. Siah2 overexpression increased the percentage of primary ciliated cells compared to that in LacZ controls. This increase was blocked by the co-expression of Par3 or PifoS (**Fig. 5C**). Taken together, these data show that primary ciliogenesis in response to Shh requires Ras or Siah2 and that Siah2 regulates primary ciliogenesis by blocking the function of Par3 and targeting the cilium disassembly protein Pifo for degradation.

### Laminin produced by the cerebellar pial epithelium supports primary ciliogenesis in GNPs in an integrin β1–Ras–dependent pathway

Having gathered evidence that the apparent regulation of Shh signaling by Ras-MAPK and Siah2 was indirect via ciliogenesis, we next sought to identify extrinsic signals that could activate Ras-MAPK in the developing EGL GZ. Ras-MAPK is classically activated by growth factors such as EGFs or IGFs and their tyrosine kinase receptors(Bergman et al., 2013; Gale et al., 1993). *In situ* data from the Brain Transcriptome Database show that the pia surrounding the EGL of a P7 cerebellum expresses insulin-like growth factor 2 (Igf2) (**fig. S10**). Given that Igf2 activates the Ras-MAPK cascade and that the high concentration of insulin in the neuronal culture supplement B27 triggers the Igf2 receptor, we wanted to rule out the possibility that Igf2 signaling controlled primary ciliogenesis and Siah2 expression. Isolated P7 GNPs from WT cerebella were placed in culture in insulin-free medium and stimulated with Igf2, Shh-N, or both in combination for 48 h (**fig. S11**). The effect of Igf2 on Siah2 expression was negligible compared to that of Shh-N, showing that Igf2 signaling plays a minimal role in regulating Siah2 expression. These data are consistent with previous reports that other receptor tyrosine kinase receptor ligands have negligible mitogenic activity for GNPs when compared to Shh(Wechsler-Reya and Scott, 1999).

GNPs express integrin β1 and interact with the laminin-rich basement membrane secreted by the pia surrounding the developing cerebellum(Blaess et al., 2004; Ichikawa-Tomikawa et al., 2012) (**Fig. 6A**). Interestingly, GNPs lacking integrin β1 are insensitive to Shh(Blaess et al., 2004), suggesting a critical role of matrix interactions in maintaining GNP responsiveness to a critical mitogen. We proceeded to place isolated GNPs from P7 cerebella in culture on monolayers of dissociated pial epithelia or glia from age-matched cerebella (**Fig. 6B**). After 48 h in culture, by which time most cells lose their primary cilia, a greater proportion of the GNPs on the pial monolayers were ciliated when compared to those on the glial monolayers. Next, we investigated whether primary ciliogenesis in GNPs on pial monolayers was dependent on integrin β1. Isolated GNPs from P7 *Integrin β1*^Flox/Flox^ cerebella were nucleofected with constructs encoding Cre recombinase or an inactive Cre mutant as a control, together with a Venus-tagged Arl13b to mark the primary cilia. The nucleofected cells were then placed in culture on pial monolayers for 48 h (**Fig. 6C**). The results showed that when integrin β1 was deleted from GNPs, fewer primary cilia were detected.

**Figure 6.**
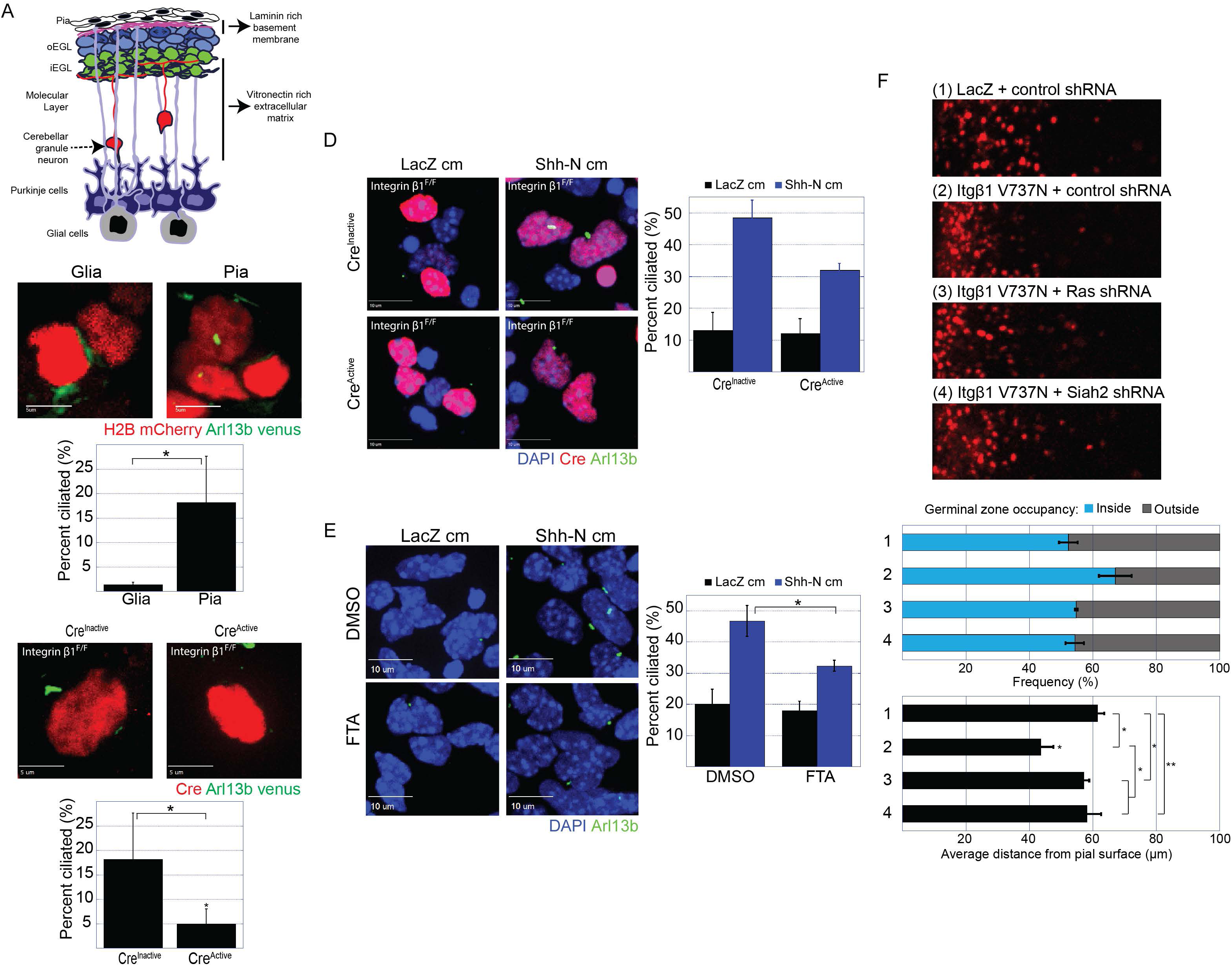
Laminin promotes primary ciliogenesis in an integrin β1-Ras–dependent pathway. (**A**) Stratification of the early postnatal cerebellum. Distinct components make up the extracellular matrix of the EGL. The cerebellar pia mater secretes a laminin-rich extracellular matrix with which proliferating GNPs make contact, whereas deeper in the cerebellum, differentiated CGNs contact a vitronectin-rich extracellular environment. The mitogenic effect of Shh is promoted by laminin but inhibited by vitronectin. (**B**) Isolated P7 CGNs were nucleofected with the bicistronic vector carrying Venus-tagged Arl13b and mCherry-tagged histone H2B and were plated on monolayers of cerebellar glial or pial cells. After 24 h in culture, the cells were fixed and processed for immunofluorescence staining followed by imaging on a spinning-disk confocal microscope. The graph shows the proportion of nucleofected cells that were ciliated. (**C–E**) P7 CGNs isolated from integrin β1–floxed cerebella were co-nucleofected with constructs expressing Venus-tagged Arl13b and a bicistronic vector carrying either the active or inactive mutant of Cre recombinase and mCherry-tagged histone H2B. The cells were cultured for 24 h on monolayers of pial cells (C) or laminin-coated slides with or without Shh-N as indicated (D and E). cm = conditioned medium. After 24 h in culture, the cells were fixed and processed for immunofluorescence staining followed by imaging on a spinning-disk confocal microscope. The proportion of nucleofected cells (H2B mCherry+) that were ciliated (C and D) or Siah2 positive (E) was determined and is shown on the accompanying graphs. (**F**) P7 cerebella were co-electroporated with H2B-mCherry and the indicated constructs. Cerebellar slices were kept in *ex vivo* culture for 48 h. Representative images of the migration patterns with the indicated manipulations are shown. The distance of H2B-labeled cells from the surface of the cerebellum was measured in three independent experiments. The graphs show the frequency distribution within and outside the EGL (top panel) and the average migration distance (lower panel). \**P* < 0.05, *\*\*P* > 0.05 (by an unpaired two-tailed *t*-test).

Having determined that the pia supported primary ciliogenesis in GNPs, we wished to determine whether this was mediated by laminin, the extracellular matrix substrate secreted by the pia. Conditional deletion of laminin from the pial basement membrane decreased proliferation in GNPs(Ichikawa-Tomikawa et al., 2012), suggesting that there was a defect in primary ciliogenesis. Accordingly, we plated isolated GNPs from P7 WT cerebella on laminin- or vitronectin-coated glass in the presence or absence of Shh-N. The latter substrate is found in the extracellular matrix surrounding the iEGL and the ML where differentiated CGNs reside and has an inhibitory effect on Shh(Pons et al., 2001). We found that laminin, but not vitronectin, supported the effect of Shh-N on both Siah2 expression **(fig. S12A)** and the primary cilia **(fig. S12B)** in GNPs. To determine whether integrin β1 was required for primary ciliogenesis when GNPs were plated on laminin, we nucleofected constructs encoding Cre recombinase, or an inactive Cre mutant as a control, into isolated GNPs from P7 *Integrin β1*^Flox/Flox^ cerebella and plated them on laminin in the presence or absence of Shh-N. When integrin β1 was deleted, fewer primary cilia were detected in response to Shh-N (**Fig. 6D**), and Siah2 expression was decreased (**Fig. S13A**). Finally, we wanted to determine whether Ras was required, as integrin β1 is known to signal to Ras. We plated isolated GNPs from P7 WT cerebella on laminin-coated glass in the presence or absence of Shh-N in combination with farnesyl thiosalicylic acid (FTA), a Ras inhibitor, or DMSO as a control. Shh-N stimulation in the presence of FTA resulted in fewer primary cilia (**fig. 6D**) and decreased Siah2 expression (**fig. S13B**).

Given that integrin β1 signaling is required for primary ciliogenesis, we wanted to know whether enhanced integrin β1 signaling would affect GZ exit in developing GNPs. To test this, we electroporated the EGLs of P7 WT cerebella with constructs encoding the auto-clustering mutant integrin β1 (V737N)(Li et al., 2003) or LacZ as a control. We found that enhanced integrin β1 signaling blocked GZ exit. We also knocked down Ras or Siah2 under the condition of enhanced integrin β1 signaling and found that GZ exit was rescued, showing that Ras or Siah2 functions downstream of integrin β1 in controlling GZ exit (**Fig. 6F**). Taken together, these data show that the pial epithelium and the laminin that it produces provide a supportive microenvironment that promotes primary ciliogenesis in GNPs, enhancing their response to Shh. Primary ciliogenesis in GNPs is mediated through integrin β1–Ras signaling.

## DISCUSSION

Throughout the developing nervous system, GZ occupancy by proliferating progenitor cells and the migration of postmitotic neurons are viewed as transient and independent phases of neuronal maturation, but there is limited insight into how these developmental stages are interconnected or potentially cross-regulated. For example, although it is apparent that radial migration is a process concomitantly linked with the cessation of progenitor proliferation and the start of neuronal terminal differentiation, it remains unknown how the cell biological programs that are necessary for migration initiation are both restrained in progenitor cells and stimulated by differentiation. It is also unclear how the cell-intrinsic machinery that organizes these processes is in turn regulated by cell-extrinsic signals, such as the mitogens that promote continued proliferation, or the contextual niche components, such as the ECM. The Siah2 E3 ubiquitin ligase appears to act as a central regulator of GZ occupancy by modulating two distinct cell biological processes. Previous results have shown that Siah2 restrains radial migration in progenitors by tagging for ubiquitin-proteasome degradation targets that are required for the pro-migratory adhesive interactions and actin-microtubule crosslinking that are essential for GZ exit and migration along glial fibers(Famulski et al., 2010; Trivedi et al., 2017). The current study has extended these findings to implicate Siah2 as an active participant in GZ occupancy by supporting primary cilium maintenance in GNPs: Siah2 targets Pard3 and Pifo for degradation, thereby ensuring the reception of signals from the Shh morphogen that promote GNP proliferation at the expense of CGN differentiation. Furthermore, the engagement of the laminin ECM substrate by integrin receptors is a parallel source of Ras-Mapk activity that is necessary for primary ciliogenesis, illustrating how coincidence detection between niche components converges on Siah2 to link GZ occupancy with proliferative signals. Intriguingly, Siah2 antagonism of Pard3 represents a mechanistic convergence point of GZ niche and mitogen coincidence detection. As Pard3 counters ciliogenesis and promotes CGN GZ exit, the relief of Siah2-target inhibition in differentiation represents a direct integration of responsiveness to a mitogen and migration initiation as GNPs mature into CGNs.

In this study, we have revealed an unexpected connection between Shh signaling, which maintains GNPs in the undifferentiated state, and Ras-Mapk and Siah2. Siah2 expression is not only maintained by Shh in a Ras-Mapk–dependent manner but is part of a functional feed-forward loop as Siah2 is in turn required for GNPs to sense an Shh signal. Although our study has not established Siah2 as a direct target of the Shh pathway, there are clear canonical aspects to Shh maintenance of Siah2 expression: 1) the gain of function associated with Shh activators such as SmoM2 and Gli1 shows that the Shh pathways are sufficient to maintain Siah2 expression; and 2) Shh-induced maintenance of Siah2 requires the primary cilia and Gli transcription factors. It is important to note that genome-wide ChIP analyses in developing GNPs have identified Siah2 as a binding target of Gli1, supporting our findings that Gli1 activates Siah2 expression(Lee et al., 2010) and that Gli transcription factors are necessary for Shh-N stimulation of Siah2 expression. Shh maintenance of Siah2 expression differs from primary Shh targets such as Ptch and Gli1 that are activated almost immediately after Shh addition(Goodrich et al., 1996; Kenney et al., 2003), because the effects of Shh on Siah2 expression required 36 to 48 h of exposure. This unique temporal variation on the usual theme of Shh-induced changes in gene expression may stem from the fact that GNPs isolated during the peak proliferative period have been pre-exposed to Shh *in vivo* and, hence, exhibit peak Siah2 expression.

Our study has broadened our knowledge of the previously reported crosstalk between Ras/Mapk and Shh signaling. Oncogenic Ras/Mapk have been demonstrated to promote the stability and transcriptional activity of Gli transcription factors in pancreatic ductal adenocarcinoma (PDAC)(Ji et al., 2007). Moreover, Ras/Mapk activity is required for Gli protein stability and nuclear translocation in melanomas(Stecca et al., 2007). In both of these cases, the loss of Ras/Mapk- or Gli1-mediated Shh signaling resulted in decreased cell proliferation similar to that which we observed in GNPs, suggesting that Shh and Ras/Mapk signaling must co-exist for optimal proliferative output. The precise mechanistic pathways regulating this crosstalk are poorly characterized in PDAC and melanoma. Our finding that Ras/Mapk support ciliogenesis suggests that maintenance of this critical Shh-transducing organelle may underlie some of the Ras/Mapk support of Shh signaling, although it is important to note that PDACs are devoid of primary cilia(Seeley et al., 2009), as is common in rapidly dividing tumors.

The process of primary ciliogenesis involves a dynamic orchestration of events, and the molecular mechanisms by which this process is regulated have been delineated using culture systems of cells that primarily elaborate cilia in a quiescent or postmitotic state(Sánchez and Dynlacht, 2016). In these models, it is thought that the primary cilium impedes cell-cycle progression and that its loss is required for re-entry into the cell cycle. A major concern with these studies, which often use serum starvation to induce primary ciliogenesis, is their lack of physiologic relevance or applicability to the mechanisms of ciliogenesis in cell types in the body that are ciliated even in the presence of active cell division. An increasing number of rapidly dividing cells or tumor cells are known to possess cilia when in the actively cycling state. The first profound illustration of this concept was the presence of primary cilia in rapidly dividing medulloblastoma tumor cells(Han et al., 2009). Our data support the shift in dogma by revealing that the primary cilia persist well into the S, G2, and early M phases of the GNP cell cycle. The structure is quickly disassembled for a short time during late M phase as a result of the process of cytokinesis, but primary cilia reappear rapidly in GNPs as they continue to proliferate if Shh is present. This suggests that the primary cilia in GNPs facilitate cell-cycle progression and, given the highly proliferative nature of these cells, that it is crucial for them to maintain their primary cilia to continuously sense Shh. We can extend this concept beyond GNP proliferation to encompass CGN migration initiation, as sustained Shh signaling and cilium loss have opposing effects on GZ exit. Thus, not only do GNPs maintain their primary cilia in order to remain sensitive to Shh, as we have shown, but differentiated CGNs retract their primary cilia, rendering the cells insensitive to Shh and enabling them to undergo radial migration.

Mechanistically, our work has revealed a novel feed-forward mechanism by which Shh signaling maintains primary ciliogenesis in proliferative GNPs. Elevated Siah2 expression in the presence of Shh supports primary ciliogenesis after GNP cell division as Siah2 is both necessary for ciliogenesis in Shh-treated GNPs and, when over-expressed, sufficient by itself to maintain cilia. How does Siah2 contribute to ciliogenesis? The substrate-binding domain of Siah ligases forms a binding groove that specifically recognizes its substrates through a degron motif: Px[ARTE]xVxP(House et al., 2003, 2006) that has remained mostly unchanged from invertebrates to humans. The high level of Siah2 expression in GNPs suggests a model whereby a group of Siah degron–containing target proteins are degraded in progenitors that probably promote cilium retraction in CGNs when Siah activity diminishes during differentiation. Our small-scale Siah2 target– expression search for proteins that could promote cilium loss revealed that Pard3 and Pifo expression in Shh-treated or Siah2-overexpressing CGNs led to cilium disassembly. In contrast, alpha-actinin 4 and drebrin, two Siah targets that are involved in uncontrolled ciliary membrane shedding in ciliopathies(Nager et al., 2017), did not induce cilium disassembly in GNPs. Although Pifo has been reported to have cilium-disassembly activities in the embryonic mouse node(Kinzel et al., 2010), the effect of Pard3 on neuronal progenitor cilia was surprising because this protein promoted cilium assembly in cells in culture that display ciliogenesis in the quiescent state(Sfakianos et al., 2007), suggesting that the mechanisms of ciliogenesis in cells that maintain their cilia while actively cycling are different from those of the classical cell types that grow cilia when quiescent. Moreover, the novel Pard complex–dependent cilium-disassembly activity and its regulation by Siah2 that was revealed by our study represents a rapid post-translational mechanism regulating GZ occupancy, whereby cilium status, polarization during differentiation, and the adhesive mechanisms are tuned via Siah2-Pard3 antagonism. Interestingly, both Pifo and Pard3, in combination with its binding partner aPKC, regulate the levels of Aurora A kinase(Khazaei and Püschel, 2009; Kinzel et al., 2010), which is a known regulator of cilium disassembly during the cell cycle, suggesting that both Pifo and Pard3 may impinge on known pathways for cilium disassembly via Aurora A(Pugacheva et al., 2007).

Previous studies demonstrated that the EGL serves as a mitogenic niche for developing GNPs and that migration away from this lamina is necessary to drive cell-cycle exit and cellular differentiation(Choi et al., 2005), hinting that the microenvironment plays a key role in controlling these transformations; however, the mechanism remains elusive. In this study, we have demonstrated that laminin, a main component of the extracellular matrix surrounding the developing EGL, supports Shh signaling in GNPs(Ichikawa-Tomikawa et al., 2012; Pons et al., 2001), and we have further determined that laminin cooperates with Shh signaling to maintain primary ciliogenesis in GNPs. Furthermore, Siah2 is a central regulator of ECM-Shh crosstalk by virtue of being downstream of Shh and Ras-Mapk, respectively, which ultimately regulates the window of GNP responsiveness to a mitogen via an unexpected function in ciliogenesis. Interestingly, laminin is present only in the basement membrane surrounding the EGL that proliferating GNPs contact(Ichikawa-Tomikawa et al., 2012; Pons et al., 2001). The laminin-rich extracellular matrix transitions to vitronectin deeper in the iEGL and ML(Hashimoto et al., 2016; Pons et al., 2001). We have provided data showing that vitronectin counters the mitogenic effect of Shh by blocking primary ciliogenesis in GNPs demonstrating an exquisite specificity of ECM to Shh coincidence detection: while both integrin β1 involved in laminin binding and integrin β3 involved in vitronectin binding both signal through the Ras-MAPK cascade only laminin support supports Shh-induced ciliogenesis. Thus, GNPs possess an ECM-mitogen coincidence detection machinery upstream of Siah2 that is sufficient to discriminate between varied Ras-Mapk activators. Such ECM-Shh coincidence detection is likely to reside outside of the nervous system as primary ciliogenesis in the developing dermal papillae epithelial cells requires integrin β1 and its interaction with laminin but is inhibited by integrin β3 (Gao et al., 2008). Given Siah2 recently expanded role in controlling epithelial polarization(Kim et al., 2014) it will be exciting to determine if ECM regulation of Siah2 activity plays a similar role in developing epithelia.

Given the pattern of migration in developing CGNs, this unique transition in components of the extracellular matrix represents a novel mechanism for controlling the fine transition between cellular proliferation and differentiation. Further studies are required to determine whether there are cell-autonomous factors that can override the cooperative effects of laminin and Shh. One interesting observation stems from the fact that most Ptch-deficient GNPs eventually exit the EGL and undergo differentiation(Kim et al., 2003). One could postulate the existence of a molecular timer in GNPs that limits their proliferative capacity and must be overcome in the event of tumorigenesis.

The newly discovered crosstalk between Shh, ECM, and Siah2 may be relevant to GZ-exit defects in cerebellar tumorigenesis. Shh-MBs arise from GNPs with aberrant Shh signaling(Dahmane and Ruiz-i-Altaba, 1999; Lewis et al., 2004; Wechsler-Reya and Scott, 1999). In humans, these tumors also exhibit increased Siah2 expression, as compared to that in other subgroups and they are ciliated. Various mouse models, and specifically the Ptch-deficient mouse model(Yang, 2008), have faithfully recapitulated this subgroup of tumors. By using a similar mouse model, we have shown that acute deletion of Ptch in GNPs blocked GZ exit, thereby recapitulating the overt phenotype seen in these preneoplastic cells. More importantly, we have shown that this phenotype is caused by the increased expression of Siah2 and that it was rescued when Siah2 was knocked down or when its function was inhibited. Overall, our data delineate a molecular mechanism that explains why GZ exit is affected in preneoplastic GNPs and why these tumor subtypes maintain their primary cilia. Given the ongoing efforts to develop Siah inhibitors(Stebbins et al., 2013), the results of this study make Siah2 an interesting potential target for treating human Shh-MB, given that reducing Siah2 activity accelerates differentiation and blocks Shh-driven primary ciliogenesis; thus, Siah inhibition has potential for use in differentiation-inducing therapies for MB.

## METHODS

### Cerebellar *ex vivo* electroporation and organotypic slice culture (*ex vivo c*erebellar pulse-chase assay)

P7 cerebellar were dissected, soaked in a suspension of plasmid DNA in Hank’s buffered salt solution (HBSS) at a concentration range of 1–3 μg/μL per construct (the nuclear marker H2B-mCherry was always included to track cellular migration), and electroporated in a platinum-block petri-dish electrode (CUY520-P5; Protech International) by using a square-wave electroporator (CUY21-EDIT; Protech International) with the following program: 5 pulses, 90 V, 50-ms pulse, 500-ms interval. The electroporated cerebella were embedded in 4% low-melting-point agarose in HBSS, and 300-μm sagittal sections of the cerebellar vermis were prepared on a vibratome (VT1200; Leica microsystems). The sections were transferred to 0.4-μm Millicell cell culture inserts (Millipore) and incubated in Basal Medium Eagle (BME) supplemented with 0.5% glucose, 2mM L-glutamine, 50 U/mL penicillin-streptomycin, and 1× B27 and 1× N2 supplements (Lifetech). After 24 or 48 h, the slices were fixed with 4% paraformaldehyde then stained with EdU, if necessary, and mounted on glass slides, with ProLong Gold antifade mountant (Lifetech) being applied before image acquisition. To measure the migration distance, the coordinates of the center of each nucleus marked by H2B-mCherry were first determined using SlideBook imaging software (3i [Intelligent Imaging Innovations]). The coordinates were then exported into the IGOR Pro line analysis software (WaveMetrics Inc.), which measures the perpendicular distance of each nucleus from the surface of the cerebellum. Statistical analyses were performed using Excel (Microsoft).

### Isolation and nucleofection of cerebellar granule neuron progenitors

P7 cerebella were dissected and dispersed using the Neural Tissue Dissociation Kit P (Miltenyi) as recommended. The tissue was titurated into a single-cell suspension by using fire-polished glass Pasteur pipettes. CGNs were isolated by centrifugation over a discontinuous Percoll gradient, and the higher-density fraction, which contained 95% CGNs and 5% glia, was collected. CGNs were nucleofected with plasmid DNA by using the optimized Amaxa Mouse Neuron Nucleofector Kit (Lonza) with the O-005 program. The cells were recovered for 5 min then plated over poly-L-ornithine-, Matrigel-, laminin-, or vitronectin-coated 16-well slides or glass-bottom dishes (MatTek Corporation) in Neurobasal medium supplemented with 0.5% glucose, 0.4 mg/mL of tissue culture– grade bovine serum albumin (Sigma), 2mM L-glutamine, 50 U/mL penicillin-streptomycin, and 1× B27 supplement (Thermo Scientific). When specified, insulin-free B27 supplement (Thermo Scientific) was used in place of the standard B27 supplement.

### Image acquisition and data analysis

Imaging was performed on a 3i Marianas Spinning Disk confocal microscope (Intelligent Imaging Innovations) consisting of a Zeiss Axio Observer microscope equipped with a 40/1.0 numerical aperture (NA) Plan-Apochromat (oil immersion) objective and a 63/1.4 NA Plan-Apochromat (oil immersion) objective. An Ultraview CSUX1 confocal head with 440–514 nm or 488–561 nm excitation filters and an ImageEM-intensified CCD camera (Hamamatsu) were used for high-resolution imaging. Images and video recordings were captured using SlideBook software (Intelligent Imaging innovations).

### Fixation and processing for 3D electron microscopy

Sagittal sections of the cerebellum (300 μm in thickness) were fixed in 2% paraformaldehyde and 2.5% glutaraldehyde in 0.1M cacodylate buffer overnight then rinsed in the same buffer. The tissue was stained using a modified heavy metal staining method(Ellisman 2010) then processed through a graded alcohol/propylene oxide series and infiltrated in propylene oxide/epon gradients. The tissue was then infiltrated overnight with 100% Epon 812 resin and polymerized for 48 h in an oven at 70°C.

### Preparation steps for focused ion beam scanning electron microscopy imaging

The sample block was mounted on an aluminum pin stub with a conductive silver paint and sputter-coated with a thin (<60 nm) layer of iridium by using a Denton DeskV sputter coater in order to electrically ground the sample and limit charging. To locate the region of interest (ROI), the block face was scanned at an acceleration voltage of 10 kV and a current of 0.2 nA on a scanning electron microscope using a concentric backscatter detector. The block was trimmed to the ROI with a Leica UC7 ultra-microtome fitted with a DiATOME knife. A relief was then milled into the block face by using a suitable ion beam current, and a protective cap was deposited using the carbon gas injection system. Fiducials were created to aid the computer vision pattern-placement algorithm. Regions of interest were imaged using the FEI Auto Slice and View software package to automate the serial-sectioning and data-collection processes.

### Focused ion beam scanning electron microscopy

The samples were imaged and processed for 3D data collection on an FEI Helios G3 system. The imaging parameters were 2 kV, 0.4 nA, and 5×5×5 nm with an in-column detector.

### Image processing and segmentation

Image stacks were aligned, filtered, cropped, and resampled to 8bit by using the FEI Amira software package.

### Western blotting

Cells were lysed in N-PER lysis buffer (Thermo Fisher) supplemented with 1× Halt Protease and Phosphatase Inhibitor Cocktail (Thermo Fisher). Lysates were reduced in Bolt Sample Reducing Agent (Thermo Fisher) and LDS loading buffer (Thermo Fisher) at 75°C for 5 min. The samples were electrophoresed on a 4%–12% SDS-PAGE gel (Thermo Fisher) and electroblotted onto a 0.45-μm Immobilon-FL polyvinylidene fluoride membrane (Millipore) by using the Mini Bolt Module wet-transfer system (Thermo Fisher). The membranes were blocked with a 1:2 dilution of Odyssey Blocking Buffer (LI-COR) in TBS for 1 h at room temperature then incubated with the following primary antibodies overnight at 4°C: anti–pan Ras provided in the Active Ras Pull-Down and Detection Kit (Cell Signaling); anti-Siah2 (Sigma, cat. no. SAB2108217); anti–β-actin (Sigma, cat. no. A2228); anti-HA (BioLegend, cat. no. MMS-101P-200); and anti-Myc (Sigma, cat. no. M4439). Odyssey IRDye secondary antibodies (LI-COR) were used for the detection on an Odyssey CLx scanner (LI-COR).

### Active Ras pull-down

Purified GNPs were pre-plated for 1 h to remove any adherent cells. Cells were collected by centrifugation at 100 × *g* for 5 min and rinsed once with ice-cold calcium- and magnesium-free PBS. They were then lysed in 1× Lysis/Binding/Wash Buffer supplemented with 1× Halt Phosphatase and Protease Inhibitor Cocktail (Thermo Fisher). Cell lysates were collected, and the protein concentration was determined using the Pierce BCA Protein Assay Kit (Thermo Scientific) and adjusted to 1 mg/mL. A total of 500 µg of total protein for each condition was treated with GTPγS or GDP at a final concentration of 10mM for 15 min at 30°C with constant agitation. The reaction was stopped by adding MgCl_2_ to a final concentration of 60mM on ice, and the reaction mixture was transferred to a spin cup with glutathione resin and 80 μg of GST-Raf1-RBD, prepared according to the manufacturer’s instructions. The reaction mixture was incubated at 4°C for 1 h with gentle rocking then washed three times with 1× Lysis/Binding/Wash Buffer. Reducing sample buffer was prepared by adding dithiothreitol to 2× SDS sample buffer to a final concentration of 200mM. Samples were eluted with 50 µL of reducing sample buffer and heated at 95°C for 5 min.

### Immunohistochemistry

*In vitro* CGN cultures were fixed with 4% paraformaldehyde for 15 min and permeabilized with 1% SDS for 20 min. This was followed by PBS washes and blocking for 30 min with 5% donkey serum. Slides were incubated overnight at 4°C with primary antibodies diluted in PBS with 0.5% donkey serum and 0.02% Triton X-100. Alexa Fluor–conjugated secondary antibodies (Lifetech) were used to detect the primary antibodies, and Prolong Gold Antifade Mountant (Lifetech) was applied to the slides before imaging.

### Tissue preparation for cryosectioning

Whole brains were dissected and fixed in 3% paraformaldehyde in PBS overnight at 4°C with gentle shaking. Specimens were then cryoprotected with 30% sucrose in PBS before being embedded in NEG-50 medium (Thermo Fisher). Sections (14 μm in thickness) were cut on a cryostat and collected on ColorFrost Plus microscope slides (Thermo Fisher). Sections were dried on a slide warmer for 30 min before immunohistochemistry staining was performed.

### Antibodies

The following antibodies were used at the dilutions specified according to whether they were used for immunohistochemistry (IHC) or Western blot analysis (WB): anti-Arl13b, 1:2000 IHC (clone N295B/66, UC Davis/NIH NeuroMab Facility), anti-Siah2, 1:200 IHC (Santa Cruz Biotechnology, cat. no. SC81787); anti-Tag1, 1:50 IHC (Developmental Studies Hybridoma Bank, cat. no. 4D7/TAG1); anti-Siah2, 1:1000 WB (Sigman, cat. no. SAB2108217); anti–pan Ras, 1:100 IHC, 1:2000 WB (Cell Signaling); anti–β-actin, 1:20000 WB (Sigma, cat. no. A2228); and anti–phospho-Mek1/2, 1:500 IHC (Cell Signaling, cat. no. 9121).

### Animals

Wildtype C57BL/6J, Ptch1^Flox/Flox^ (B6N.129-*Ptch1^tm1Hahn^*/J) and Integrin β1^Flox/ Flox^ mice (B6;129-*Itgb1^tm1Efu^*/J) were obtained from The Jackson Laboratory. All mouse lines were maintained in standard conditions in accordance with guidelines approved by the Institutional Animal Care and Use Committee at St. Jude Children’s Research Hospital (protocol no. 483).

## ACKNOWLEDGEMENTS

We thank the Cell and Tissue Imaging Center of St. Jude Children’s Research Hospital for assistance implementing imaging analysis support and Yohann Garnier for performing preliminary forms of some of the Ras inhibitor experiments. Young Goo Han, PhD provided Ift88 shRNA reagents. Keith A. Laycock, PhD, ELS edited the manuscript. The Solecki Laboratory is funded by the American Lebanese Syrian Associated Charities (ALSAC) and by grants 1R01NS066936 and R01NS104029-02 from the National Institute of Neurological Disorders (NINDS). The content is solely the responsibility of the authors and does not necessarily represent the official views of the NINDS or the NIH.

## Author Contributions

TO carried out all *in vitro*/*ex vivo* studies, Siah/Ptch/Ras/Mapk epistasis analyses, prepared all figures. NT carried out pial co-culture experiments. RW and SF spearheaded LLS Helios FIB-SEM imaging. DJS conceived of the study, participated in its design and coordination and performed preliminary versions of some *ex vivo* experiments. All authors drafted or edited the manuscript.

## Supplementary Figure Legends

**figure S1.**
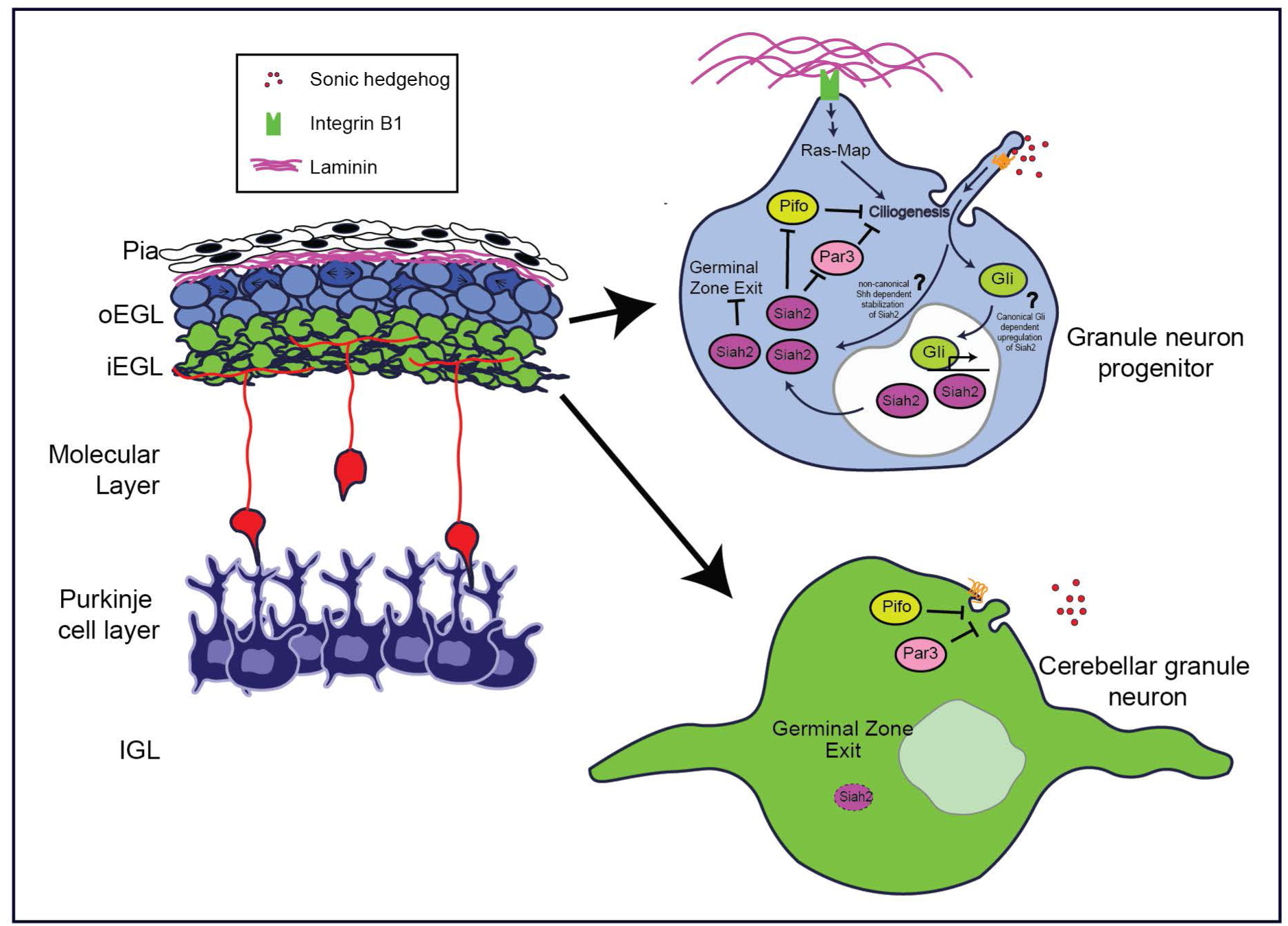
Schematic of how Siah2 is regulated in developing CGNs. The laminin rich microenvironment of the oEGL promotes primary ciliogenesis in GNPs through Integrin β1 – Ras/Mapk signaling, which allows them to sense the mitogen Shh. Activation of the Shh pathway maintains Siah2 expression which in turn promotes GZ occupancy by inhibiting GZ exit and maintaining mitogen sensitivity by promoting primary ciliogenesis through the antagonism of a key cilia disassembly protein Pifo and the polarity inducer Par3. As GNPs leave the oEGL, the lack of trophic support leads to the disassembly of the primary cilia and loss of sensitivity to Shh, which promotes CGN differentiation.

**figure S2.**
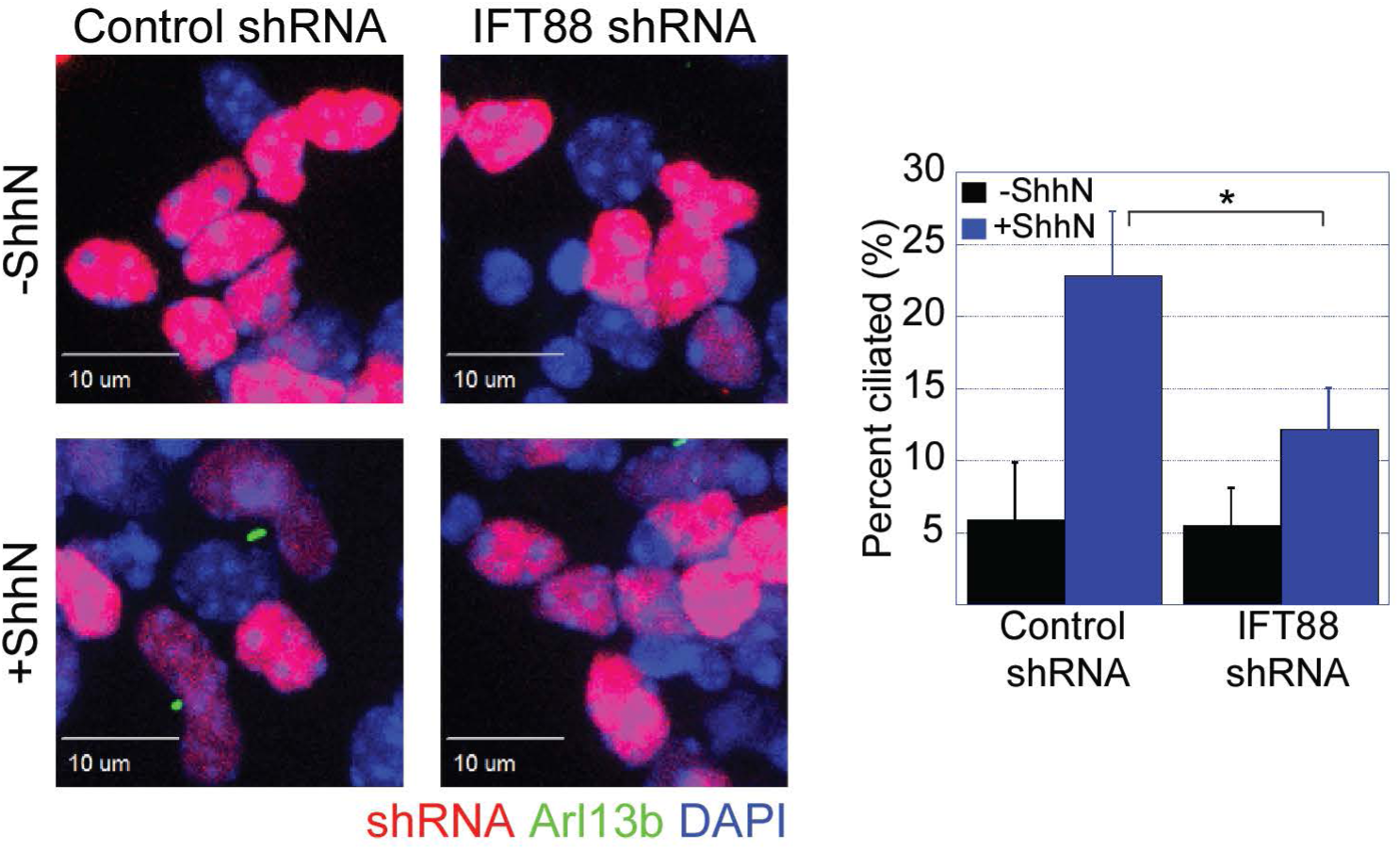
Knockdown of IFT88 impairs Shh-driven primary ciliogenesis. Isolated P7 CGNs were co-nucleofected with the nuclear marker H2B-mCherry and IFT88 shRNA and cultured *in vitro* for 48 h in the presence or absence of Shh-N (3μg/mL). The cells were then fixed, and immunofluorescence staining for Arl13b was performed. The cells were then imaged on a spinning-disk confocal microscope. The graph shows the percentage of ciliated cells, as determined by manual quantification. \**P* < 0.05 (by unpaired two-tailed *t*-test).

**figure S3.**
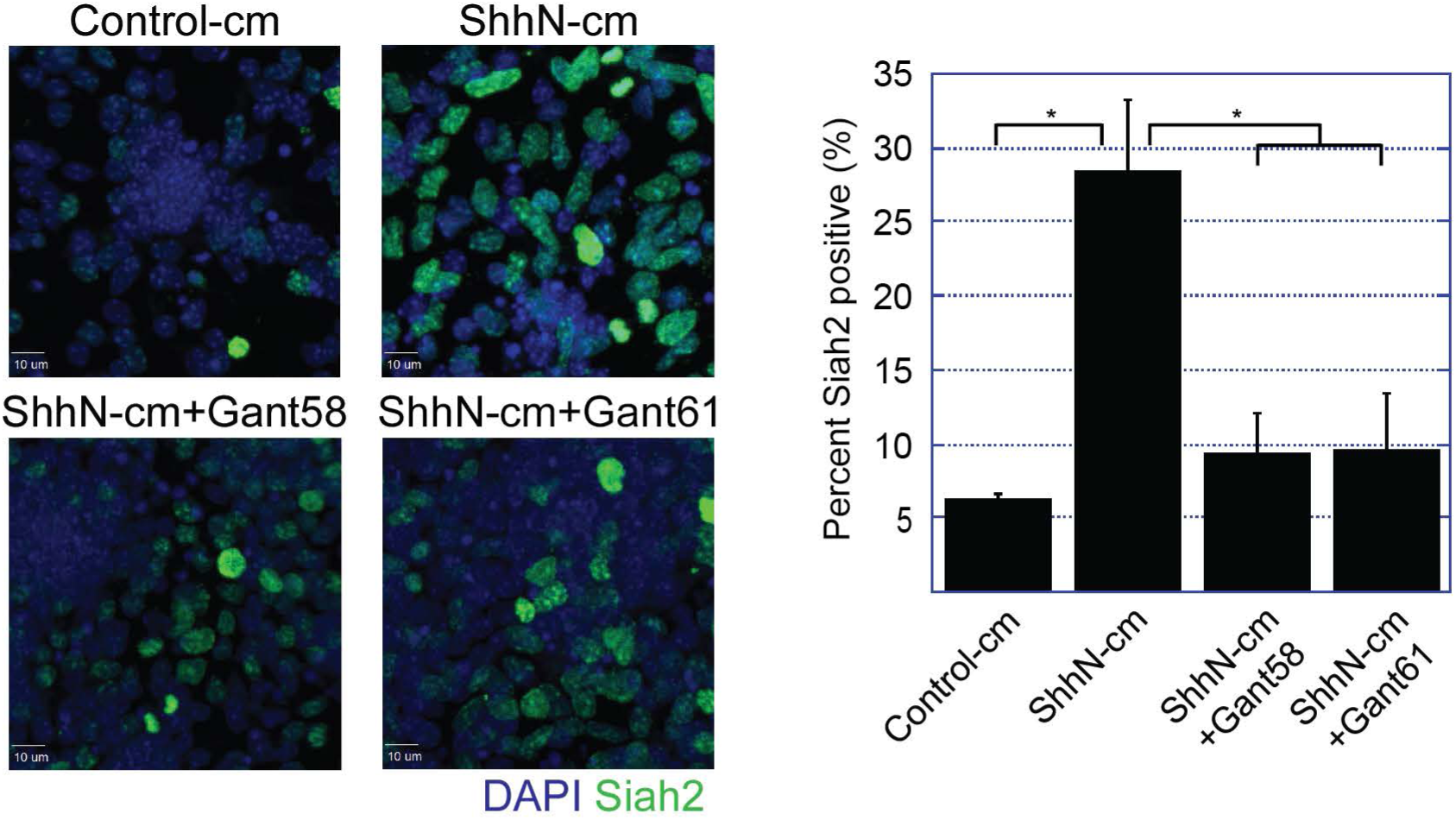
Shh maintenance of Siah2 requires Gli1 and Gli2. Isolated P7 CGNs were plated on matrigel-coated slides as indicated in combination with a Gli1 inhibitor, Gant58, or Gli1 and Gli2 inhibitor, Gant61, for 48 h. cm = conditioned medium. The cells were then fixed, and immunofluorescence staining for Siah2 was performed. The cells were then imaged on a spinning-disk confocal microscope. The graph shows the percentage of Siah2 positive cells, as determined using Slidebook. \**P* < 0.05 (by unpaired two-tailed *t*-test).

**figure S4.**
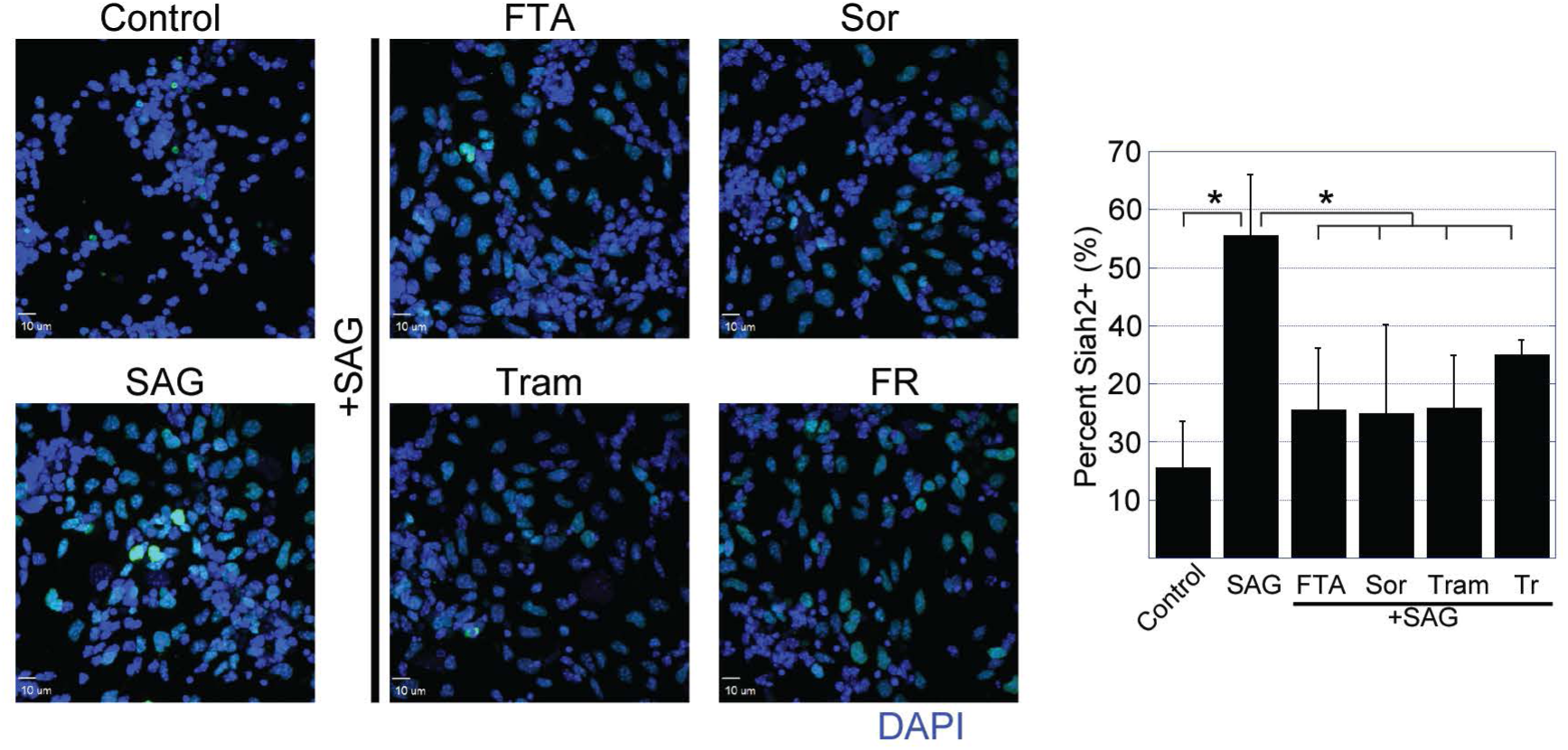
Shh maintenance of Siah2 requires the Ras/Mapk cascade. Isolated P7 CGNs were cultured for 48 h in the presence or absence of Shh agonist (SAG, 100nM) and with the indicated combinations of Ras/Mapk inhibitors: farnesyl transferase inhibitor (FTA), sorafenib (Sor), trametinib (Tram), and FR180204 (FR), which inhibit Ras, Raf, Mek1/2, and Erk1/2, respectively. The cells were fixed, and immunofluorescence staining for Siah2 was performed. The percentage of Siah2-positive cells was determined using SlideBook, and the graph shows the results. \**P* < 0.05 (by an unpaired two-tailed *t*-test).

**figure S5.**
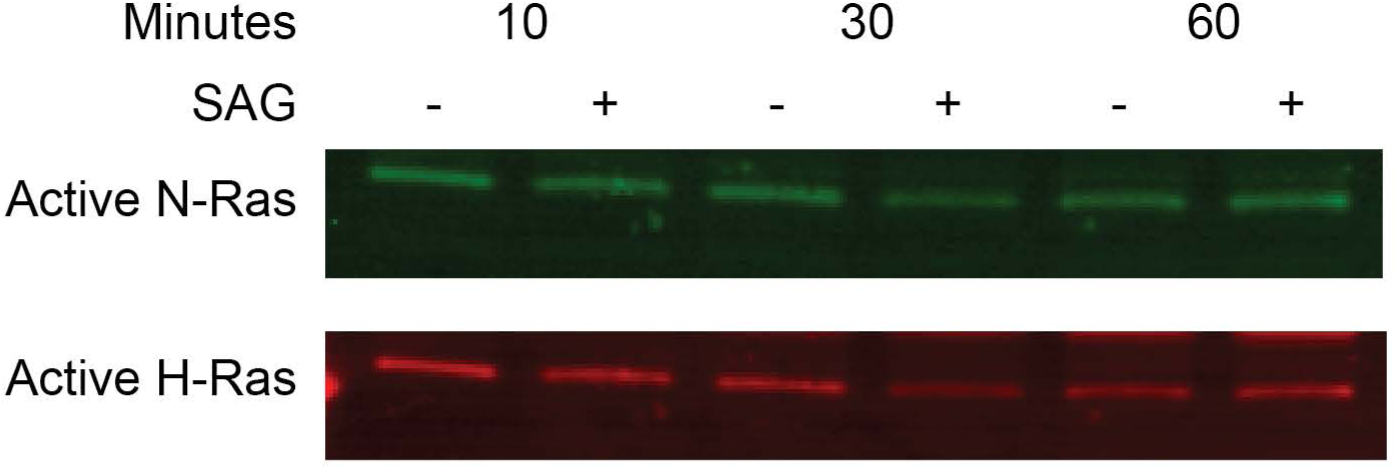
Shh pathway stimulation in GNPs does not activate Ras. Isolated P7 CGNs were plated and allowed to rest for 3 h in serum-free medium before being stimulated with SAG (100nM) at the indicated time-points. Lysates were collected and processed for active Ras pull-down by using the Active Ras Detection Kit (Cell Signaling) in accordance with the manufacturer’s instructions. Western blot analysis for H- and N-Ras was then performed.

**figure S6.**
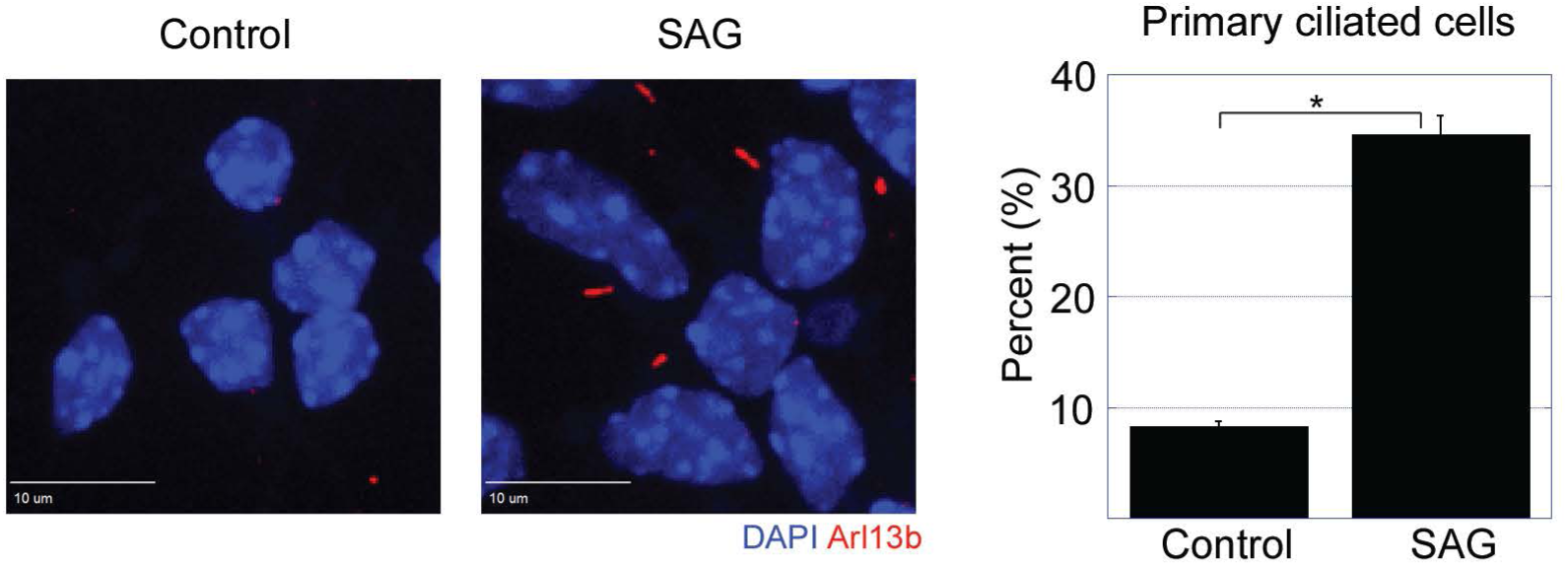
Shh signaling promotes primary ciliogenesis in GNPs. Isolated P7 CGNs were cultured in the presence or absence of SAG (100nM) for 48 h. Cells were fixed, and immunofluorescence staining for Arl13b was performed, followed by imaging with a spinning-disk confocal microscope. The graph shows the percentage of ciliated cells, as determined by manual scoring. \**P* < 0.05 (by an unpaired two-tailed *t*-test).

**figure S7.**
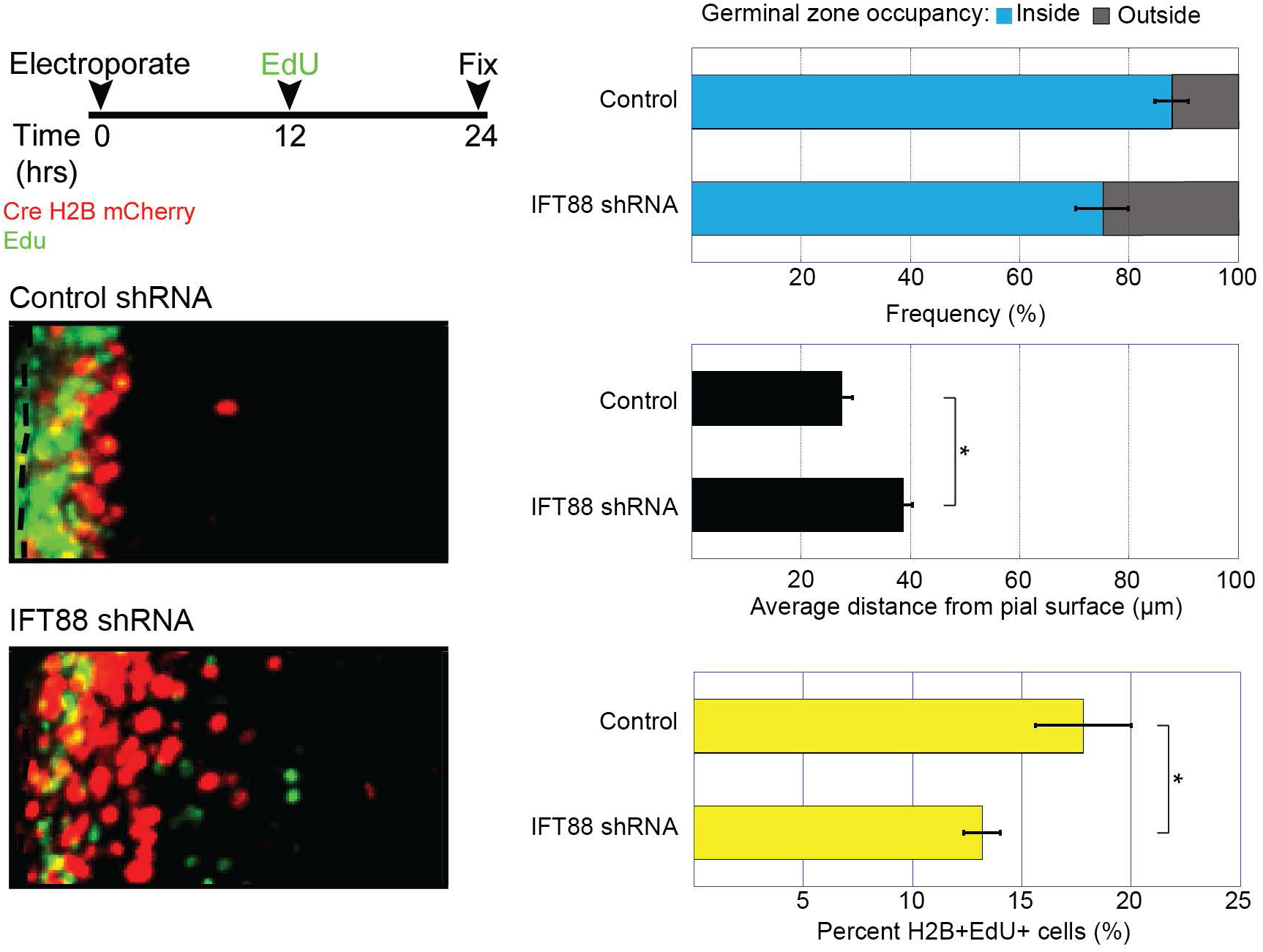
Knockdown of IFT88 promotes cell-cycle exit and spurs early GZ exit. P7 cerebella were co-electroporated with H2B-mCherry and the indicated shRNAs. Cerebellar slices were prepared and kept in *ex vivo* culture for 24 h with the addition of EdU for the final 12 h of culture. The slices were then fixed, EdU staining was performed, and the slices were imaged on a spinning-disk confocal microscope. Representative images of the migration patterns with the indicated manipulations are shown. The distance of H2B-labeled cells from the surface of the cerebellum was measured in three independent experiments. The graphs show the migration distribution, average distances, and the results of proliferative index analyses. \**P* < 0.05 (by an unpaired two-tailed *t-*test).

**figure S8.**
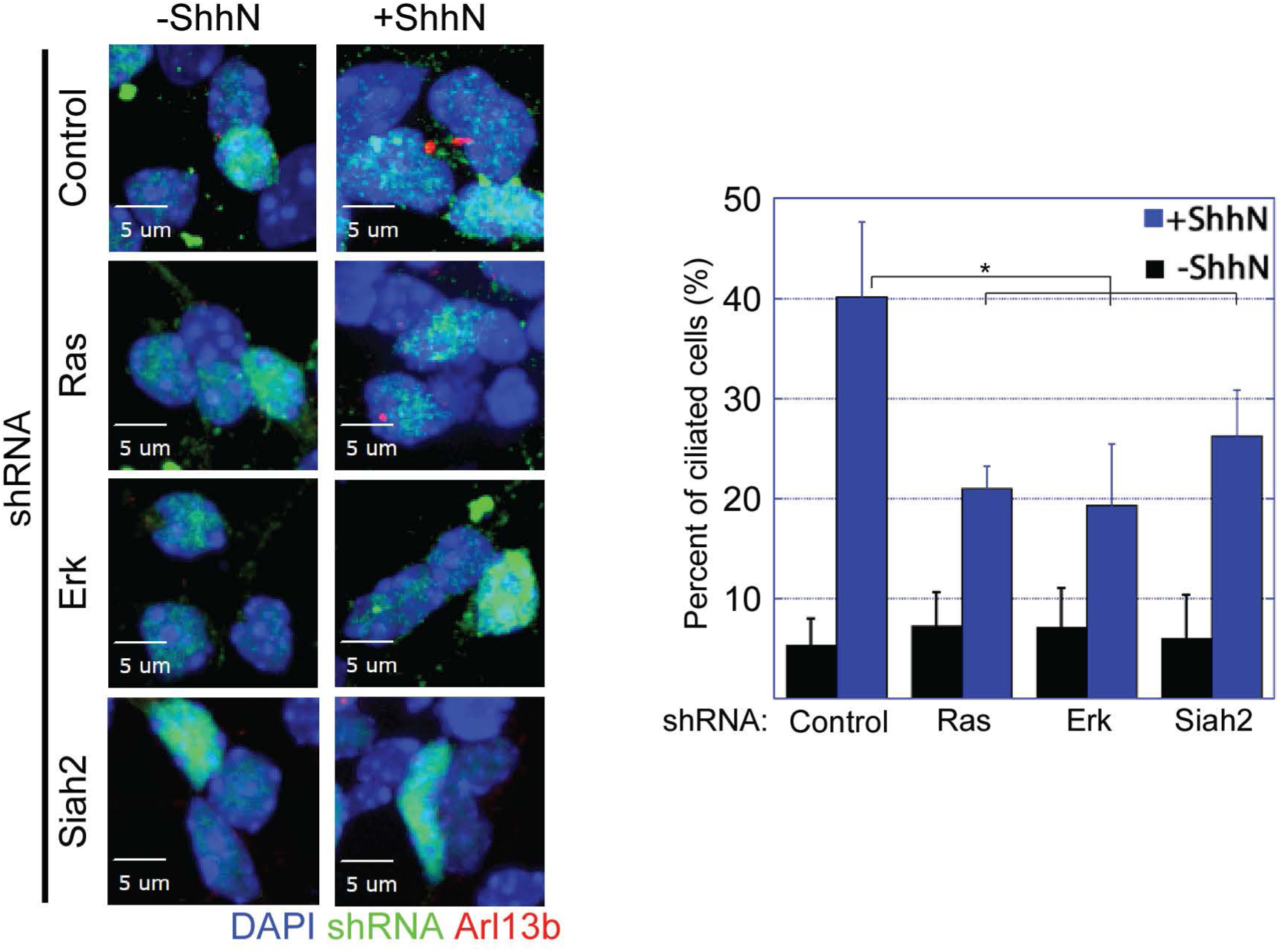
Shh-driven primary ciliogenesis requires Ras, Erk1/2, and Siah2. Isolated P7 CGNs were nucleofected with the indicated shRNAs and cultured for 48 h in the presence or absence of Shh-N (3 μg/mL). The cells were then fixed, and immunofluorescence staining for Arl13b was performed, followed by imaging on a spinning-disk confocal microscope. The graph shows the percentage of nucleofected cells that were ciliated, as determined by manual scoring. \**P* < 0.05 (by an unpaired two-tailed *t-*test).

**figure S9.**
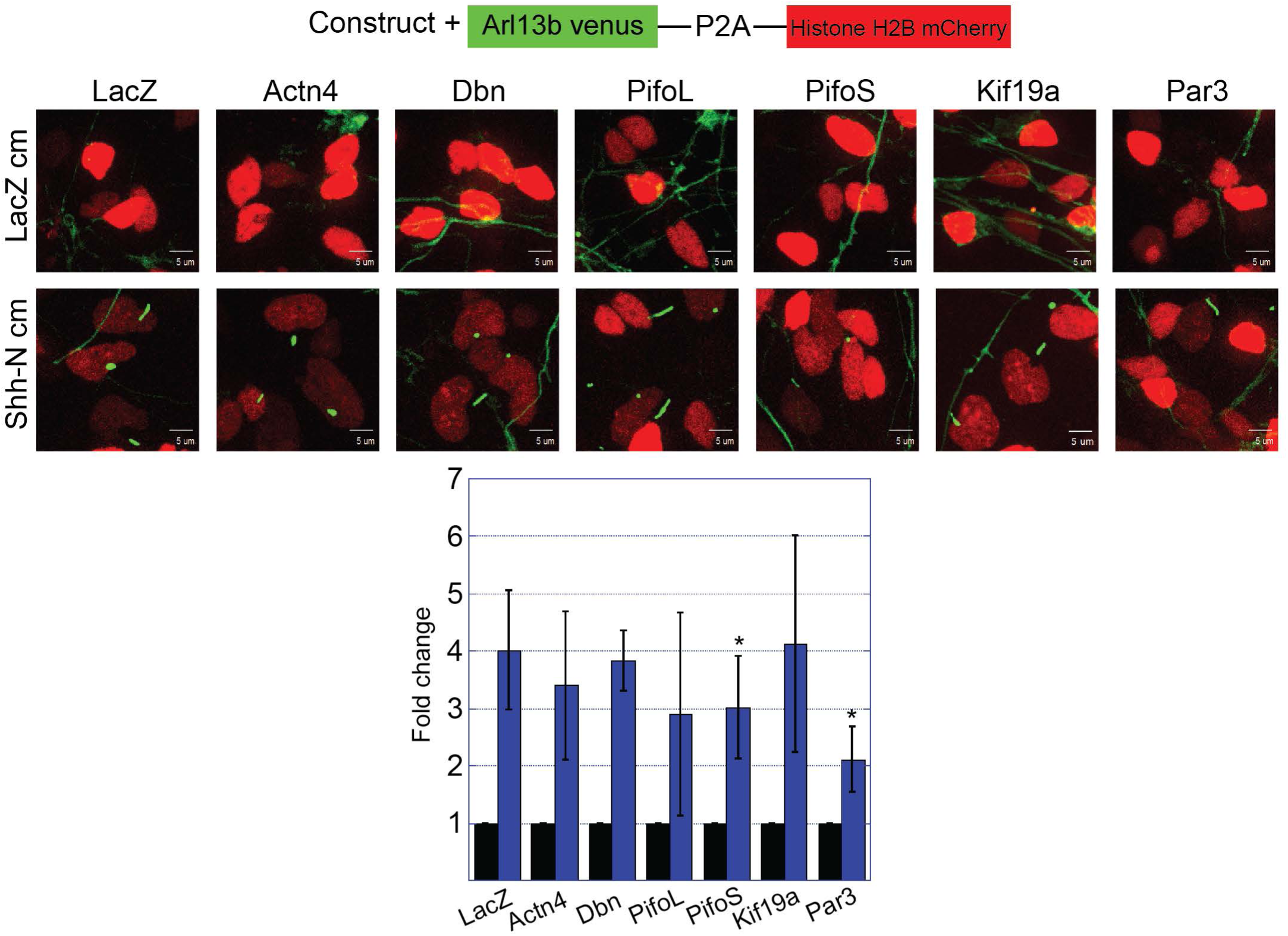
Overexpression of Pifo and Par3 blocks Shh-driven primary ciliogenesis. Isolated P7 CGNs were co-nucleofected with the indicated constructs and a bicistronic vector encoding Venus-tagged Arl13b and mCherry-tagged histone H2B. The cells were cultured overnight in the presence or absence of Shh-N–conditioned medium. Live-cell imaging with a spinning-disk confocal microscope was performed, and the percentage of nucleofected cells (H2B mCherry+) that were ciliated was determined by manual scoring, as shown on the graph. \**P* < 0.05 (by an unpaired two-tailed *t-*test).

**figure S10.**
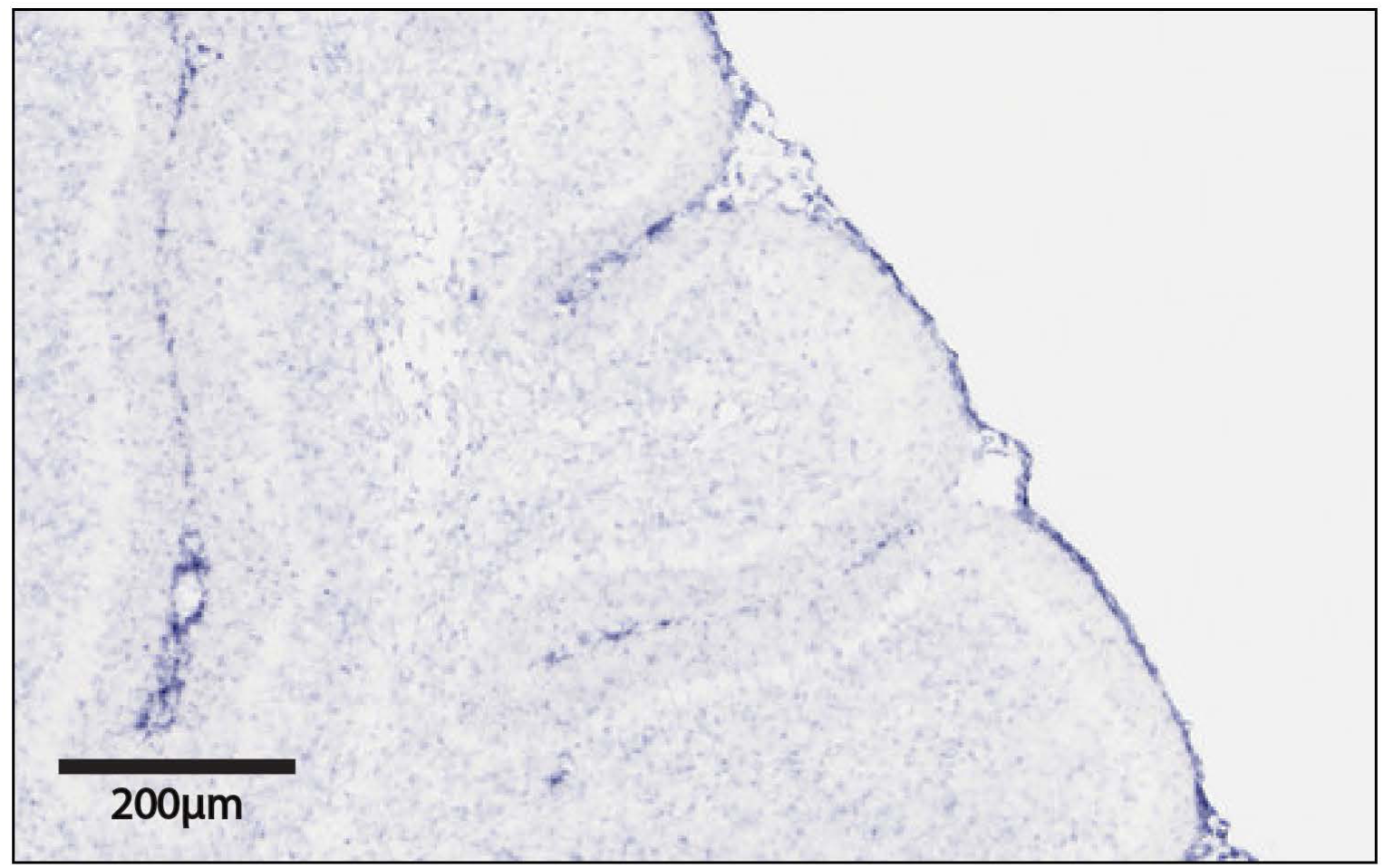
The pia mater surrounding the P7 cerebellum produces IGF2. Results of *in situ* hybridization analysis of IGF2 mRNA in a P7 cerebellum. Data were obtained from the Brain Transcriptome Database (BrainTx).

**figure S11.**
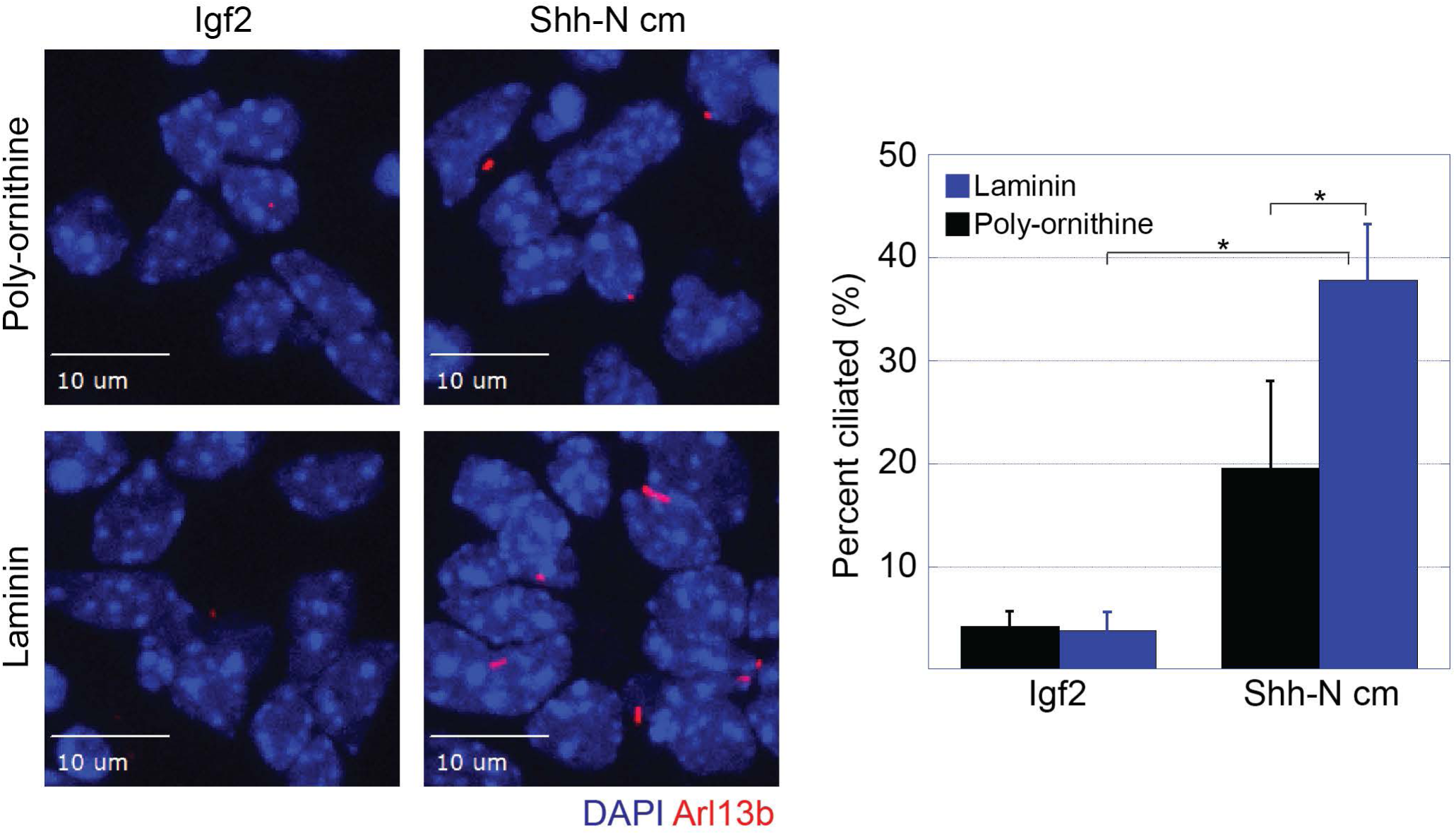
Laminin promotes Shh-driven primary ciliogenesis. Isolated P7 CGNs were plated on poly-ornithine– or laminin-coated slides and cultured in either Igf2 (200 ng/mL)- or Shh-N–conditioned medium for 48 h. The cells were then fixed, and immunofluorescence staining for Arl13b was performed. The cells were then imaged on a spinning-disk confocal microscope. The graph shows the percentage of ciliated cells, as determined by manual scoring. \**P* < 0.05, \*\**P* > 0.05 (by an unpaired two-tailed *t-* test).

**figure S12.**
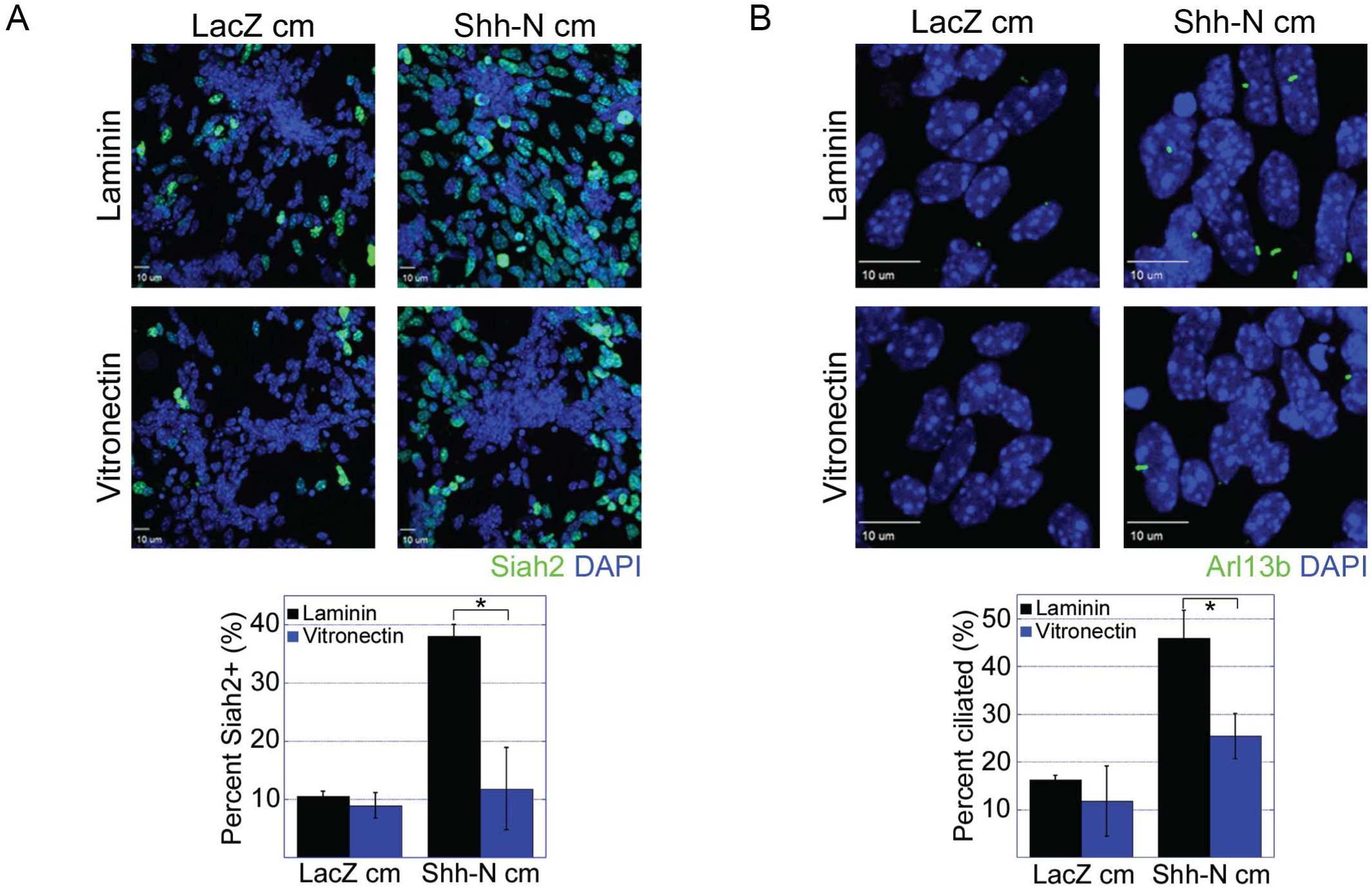
Laminin promotes whereas vitronectin inhibits Shh-driven primary ciliogenesis and Siah2 expression. Isolated P7 CGNs were plated on either laminin- or vitronectin-coated slides for 48 h in the indicated conditioned media. cm = conditioned medium. The cells were then fixed, and immunofluorescence staining for Siah2 (**A**) or Arl13b (**B**) was performed. Images were acquired on a spinning-disk confocal microscope, and the percentage of Siah2-positive cells was determined using SlideBook (A) or the percentage of ciliated cells was determined by manual scoring (B). The graphs show the results of these quantifications. \**P* < 0.05 (by an unpaired two-tailed *t*-test).

**figure S13.**
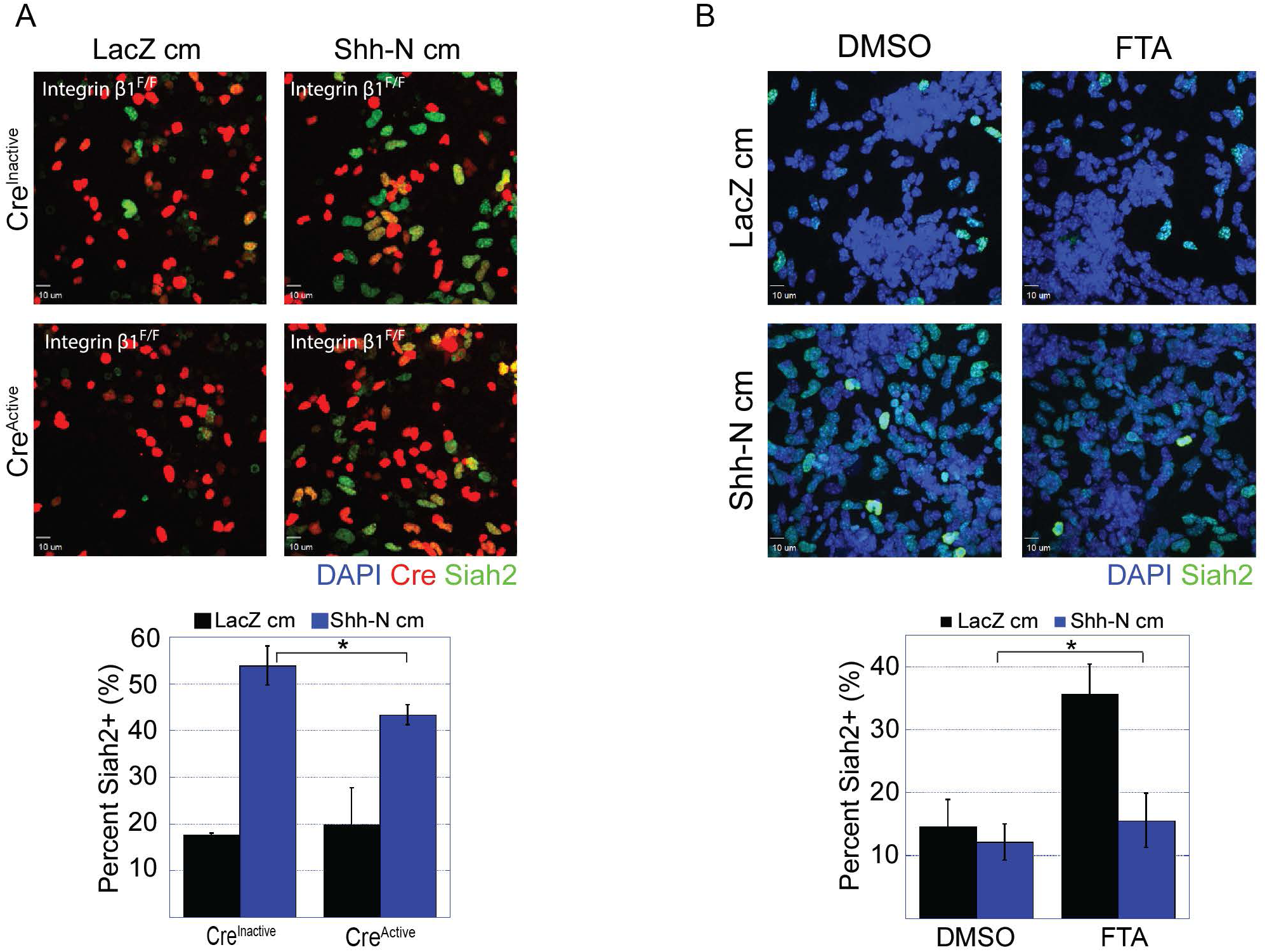
Laminin promotes whereas vitronectin inhibits Shh-driven primary ciliogenesis and Siah2 expression. Isolated P7 CGNs were plated on laminin-coated slides for 48 h as indicated. cm = conditioned medium, DMSO = dimethyl sulfoxide, FTA = farnesyl thiosalicylic acid. The cells were then fixed, and immunofluorescence staining for Arl13b (**A**) or Siah2 (**B**) was performed. Images were acquired on a spinning-disk confocal microscope, and the percentage of ciliated cells was determined by manual scoring (A) or the percentage of Siah2-positive cells was determined using SlideBook (B). The graphs show the results of these quantifications. \**P* < 0.01 (by an unpaired two-tailed *t*-test).

## REFERENCES

Almodovar, C.R. de, Coulon, C., Salin, P.A., Knevels, E., Chounlamountri, N., Poesen, K., Hermans, K., Lambrechts, D., Geyte, K.V., Dhondt, J., et al. (2010). Matrix-Binding Vascular Endothelial Growth Factor (VEGF) Isoforms Guide Granule Cell Migration in the Cerebellum via VEGF Receptor Flk1. J. Neurosci. 30, 15052–15066.

Aruga, J., Inoue, T., Hoshino, J., and Mikoshiba, K. (2002). Zic2 Controls Cerebellar Development in Cooperation with Zic1. J. Neurosci. 22, 218–225.

Bar-Sagi, D., and Hall, A. (2000). Ras and Rho GTPases: A Family Reunion. Cell 103, 227–238.

Bergman, D., Halje, M., Nordin, M., and Engström, W. (2013). Insulin-Like Growth Factor 2 in Development and Disease: A Mini-Review. Gerontology 59, 240–249.

Bishop, G.A., Berbari, N.F., Lewis, J., and Mykytyn, K. (2007). Type III adenylyl cyclase localizes to primary cilia throughout the adult mouse brain. J. Comp. Neurol. 505, 562–571.

Blaess, S., Graus-Porta, D., Belvindrah, R., Radakovits, R., Pons, S., Littlewood-Evans, A., Senften, M., Guo, H., Li, Y., Miner, J.H., et al. (2004). β1-Integrins Are Critical for Cerebellar Granule Cell Precursor Proliferation. J. Neurosci. 24, 3402–3412.

Butts, T., Green, M.J., and Wingate, R.J.T. (2014). Development of the cerebellum: simple steps to make a ‘little brain.’ Development 141, 4031–4041.

Capon, D.J., Seeburg, P.H., McGrath, J.P., Hayflick, J.S., Edman, U., Levinson, A.D., and Goeddel, D.V. (1983). Activation of Ki- ras 2 gene in human colon and lung carcinomas by two different point mutations. Nature 304, 507.

Carthew, R.W., and Rubin, G.M. (1990). seven in absentia, a gene required for specification of R7 cell fate in the Drosophila eye. Cell 63, 561–577.

Caspary, T., Larkins, C.E., and Anderson, K.V. (2007). The Graded Response to Sonic Hedgehog Depends on Cilia Architecture. Dev. Cell 12, 767–778.

Chen, C.-H., von Kessler, D.P., Park, W., Wang, B., Ma, Y., and Beachy, P.A. (1999). Nuclear Trafficking of Cubitus interruptus in the Transcriptional Regulation of Hedgehog Target Gene Expression. Cell 98, 305–316.

Chizhikov VV, Davenport J, Zhang Q, Shih EK, Cabello OA, Fuchs JL, Yoder BK, Millen K. (2007). Cilia proteins control cerebellar morphogenesis by promoting expansion of the granule progenitor pool. Journal of Neuroscience 27(36):9780–9

Choi, Y., Borghesani, P.R., Chan, J.A., and Segal, R.A. (2005). Migration from a Mitogenic Niche Promotes Cell-Cycle Exit. J. Neurosci. 25, 10437–10445.

Cox, A.D., and Der, C.J. (2010). Ras history. Small GTPases 1, 2–27.

Dahmane, N., and Ruiz-i-Altaba, A. (1999). Sonic hedgehog regulates the growth and patterning of the cerebellum. Development 126, 3089–3100.

Famulski, J.K., Trivedi, N., Howell, D., Yang, Y., Tong, Y., Gilbertson, R., and Solecki, D.J. (2010). Siah Regulation of Pard3A Controls Neuronal Cell Adhesion During Germinal Zone Exit. Science 330, 1834–1838.

Fortini, M.E., Simon, M.A., and Rubin, G.M. (1992). Signalling by the sevenless protein tyrosine kinase is mimicked by Rasl activation. Nature 355, 559–561.

Gale, N.W., Kaplan, S., Lowenstein, E.J., Schlessinger, J., and Bar-Sagi, D. (1993). Grb2 mediates the EGF-dependent activation of guanine nucleotide exchange on Ras. Nature 363, 88.

Gao, J., DeRouen, M.C., Chen, C.-H., Nguyen, M., Nguyen, N.T., Ido, H., Harada, K., Sekiguchi, K., Morgan, B.A., Miner, J.H., et al. (2008). Laminin-511 is an epithelial message promoting dermal papilla development and function during early hair morphogenesis. Genes Dev. 22, 2111–2124.

Gao, W.-Q., Heintz, N., and Hatten, M.E. (1991). Cerebellar granule cell neurogenesis is regulated by cell-cell interactions in vitro. Neuron 6, 705–715.

Gilbertson, R.J. (2008). The origins of medulloblastoma subtypes. Annu Rev Pathol 3, 341–365.

Goodrich, L.V., Johnson, R.L., Milenkovic, L., McMahon, J.A., and Scott, M.P. (1996). Conservation of the hedgehog/patched signaling pathway from flies to mice: induction of a mouse patched gene by Hedgehog. Genes Dev. 10, 301–312.

Goodrich, L.V., Milenković, L., Higgins, K.M., and Scott, M.P. (1997). Altered Neural Cell Fates and Medulloblastoma in Mouse patched Mutants. Science 277, 1109–1113.

Graus-Porta, D., Blaess, S., Senften, M., Littlewood-Evans, A., Damsky, C., Huang, Z., Orban, P., Klein, R., Schittny, J.C., and Müller, U. (2001). β1-Class Integrins Regulate the Development of Laminae and Folia in the Cerebral and Cerebellar Cortex. Neuron 31, 367–379.

Han, Y.-G., Kim, H.J., Dlugosz, A.A., Ellison, D.W., Gilbertson, R.J., and Alvarez-Buylla, A. (2009). Dual and opposing roles of primary cilia in medulloblastoma development. Nat. Med. 15, 1062–1065.

Hashimoto, K., Sakane, F., Ikeda, N., Akiyama, A., Sugahara, M., and Miyamoto, Y. (2016). Vitronectin promotes the progress of the initial differentiation stage in cerebellar granule cells. Mol. Cell. Neurosci. 70, 76–85.

Hatten, M.E. (1999). Central Nervous System Neuronal Migration. Annu. Rev. Neurosci. 22, 511–539.

Hatten, M.E. and Roussel, M.F. (2011). Development and Cancer of the Cerebellum. Trends in Neurosci. 34(3):134–42.

House, C.M., Frew, I.J., Huang, H.-L., Wiche, G., Traficante, N., Nice, E., Catimel, B., and Bowtell, D.D.L. (2003). A binding motif for Siah ubiquitin ligase. Proc. Natl. Acad. Sci. 100, 3101–3106.

House, C.M., Hancock, N.C., Möller, A., Cromer, B.A., Fedorov, V., Bowtell, D.D.L., Parker, M.W., and Polekhina, G. (2006). Elucidation of the Substrate Binding Site of Siah Ubiquitin Ligase. Structure 14, 695–701.

Hu, G., and Fearon, E.R. (1999). Siah-1 N-Terminal RING Domain Is Required for Proteolysis Function, and C-Terminal Sequences Regulate Oligomerization and Binding to Target Proteins. Mol. Cell. Biol. 19, 724–732.

Huangfu, D., and Anderson, K.V. (2005). Cilia and Hedgehog responsiveness in the mouse. Proc. Natl. Acad. Sci. U. S. A. 102, 11325–11330.

Huangfu, D., Liu, A., Rakeman, A.S., Murcia, N.S., Niswander, L., and Anderson, K.V. (2003). Hedgehog signalling in the mouse requires intraflagellar transport proteins. Nature 426, 83.

Ichikawa-Tomikawa, N., Ogawa, J., Douet, V., Xu, Z., Kamikubo, Y., Sakurai, T., Kohsaka, S., Chiba, H., Hattori, N., Yamada, Y., et al. (2012). Laminin α1 is essential for mouse cerebellar development. Matrix Biol. 31, 17–28.

Ji, Z., Mei, F.C., Xie, J., and Cheng, X. (2007). Oncogenic KRAS Activates Hedgehog Signaling Pathway in Pancreatic Cancer Cells. J. Biol. Chem. 282, 14048–14055.

Kenney, A.M., Cole, M.D., and Rowitch, D.H. (2003). *Nmyc* upregulation by sonic hedgehog signaling promotes proliferation in developing cerebellar granule neuron precursors. Development 130, 15–28.

Khazaei, M.R., and Püschel, A.W. (2009). Phosphorylation of the Par Polarity Complex Protein Par3 at Serine 962 Is Mediated by Aurora A and Regulates Its Function in Neuronal Polarity. J. Biol. Chem. 284, 33571–33579.

Kim, H., Claps, G., Möller, A., Bowtell, D., Lu, X., and Ronai, Z.A. (2014). Siah2 regulates tight junction integrity and cell polarity through control of ASPP2 stability. Oncogene 33, 2004–2010.

Kim, J.Y.H., Nelson, A.L., Algon, S.A., Graves, O., Sturla, L.M., Goumnerova, L.C., Rowitch, D.H., Segal, R.A., and Pomeroy, S.L. (2003). Medulloblastoma tumorigenesis diverges from cerebellar granule cell differentiation in patched heterozygous mice. Dev. Biol. 263, 50–66.

Kinzel, D., Boldt, K., Davis, E.E., Burtscher, I., Trümbach, D., Diplas, B., Attié-Bitach, T., Wurst, W., Katsanis, N., Ueffing, M., et al. (2010). Pitchfork Regulates Primary Cilia Disassembly and Left-Right Asymmetry. Dev. Cell 19, 66–77.

Lee, E.Y., Ji, H., Ouyang, Z., Zhou, B., Ma, W., Vokes, S.A., McMahon, A.P., Wong, W.H., and Scott, M.P. (2010). Hedgehog pathway-regulated gene networks in cerebellum development and tumorigenesis. Proc. Natl. Acad. Sci. 107, 9736–9741.

Lewis, P.M., Gritli-Linde, A., Smeyne, R., Kottmann, A., and McMahon, A.P. (2004). Sonic hedgehog signaling is required for expansion of granule neuron precursors and patterning of the mouse cerebellum. Dev. Biol. 270, 393–410.

Li, R., Mitra, N., Gratkowski, H., Vilaire, G., Litvinov, R., Nagasami, C., Weisel, J.W., Lear, J.D., DeGrado, W.F., and Bennett, J.S. (2003). Activation of Integrin αIIbß3 by Modulation of Transmembrane Helix Associations. Science 300, 795–798.

McCubrey, J.A., Steelman, L.S., Chappell, W.H., Abrams, S.L., Wong, E.W.T., Chang, F., Lehmann, B., Terrian, D.M., Milella, M., Tafuri, A., et al. (2007). Roles of the Raf/MEK/ERK pathway in cell growth, malignant transformation and drug resistance. Biochim. Biophys. Acta BBA - Mol. Cell Res. 1773, 1263–1284.

Miyata, T., Maeda, T., and Lee, J.E. (1999). NeuroD is required for differentiation of the granule cells in the cerebellum and hippocampus. Genes Dev. 13, 1647–1652.

Nager, A.R., Goldstein, J.S., Herranz-Pérez, V., Portran, D., Ye, F., Garcia-Verdugo, J.M., and Nachury, M.V. (2017). An Actin Network Dispatches Ciliary GPCRs into Extracellular Vesicles to Modulate Signaling. Cell 168, 252–263.e14.

Niwa, S., Nakajima, K., Miki, H., Minato, Y., Wang, D., and Hirokawa, N. (2012). KIF19A Is a Microtubule-Depolymerizing Kinesin for Ciliary Length Control. Dev. Cell 23, 1167–1175.

Oliver, T.G., Read, T.A., Kessler, J.D., Mehmeti, A., Wells, J.F., Huynh, T.T.T., Lin, S.M., and Wechsler-Reya, R.J. (2005). Loss of *patched* and disruption of granule cell development in a pre-neoplastic stage of medulloblastoma. Development 132, 2425–2439.

Pazour, G.J., Dickert, B.L., Vucica, Y., Seeley, E.S., Rosenbaum, J.L., Witman, G.B., and Cole, D.G. (2000). Chlamydomonas IFT88 and Its Mouse Homologue, Polycystic Kidney Disease Gene Tg737, Are Required for Assembly of Cilia and Flagella. J. Cell Biol. 151, 709–718.

Penas, C., Govek, E.-E., Fang, Y., Ramachandran, V., Daniel, M., Wang, W., Maloof, M.E., Rahaim, R.J., Bibian, M., Kawauchi, D., et al. (2015). Casein Kinase 1δ Is an APC/CCdh1 Substrate that Regulates Cerebellar Granule Cell Neurogenesis. Cell Rep. 11, 249–260.

Pickford, L.B., Mayer, D.N., Bolin, L.M., and Rouse, R.V. (1989). Transiently expressed, neural-specific molecule associated with premigratory granule cells in postnatal mouse cerebellum. J. Neurocytol. 18, 465–478.

Pons, S., Trejo, J.L., Martinez-Morales, J.R., and Marti, E. (2001). Vitronectin regulates Sonic hedgehog activity during cerebellum development through CREB phosphorylation. Development 128, 1481–1492.

Pugacheva, E.N., Jablonski, S.A., Hartman, T.R., Henske, E.P., and Golemis, E.A. (2007). HEF1-Dependent Aurora A Activation Induces Disassembly of the Primary Cilium. Cell 129, 1351–1363.

Qi, X., Schmiege, P., Coutavas, E., Wang, J., and Li, X. (2018). Structures of human Patched and its complex with native palmitoylated sonic hedgehog. Nature 560, 128.

Roessler, E., Ermilov, A.N., Grange, D.K., Wang, A., Grachtchouk, M., Dlugosz, A.A., and Muenke, M. (2005). A previously unidentified amino-terminal domain regulates transcriptional activity of wild-type and disease-associated human GLI2. Hum. Mol. Genet. 14, 2181–2188.

Rohatgi, R. (2007). Patched1 regulates hedgehog signaling at the primary cilium. Science 317, 372–376.

Sakaue-Sawano, A., Kurokawa, H., Morimura, T., Hanyu, A., Hama, H., Osawa, H., Kashiwagi, S., Fukami, K., Miyata, T., Miyoshi, H., et al. (2008). Visualizing Spatiotemporal Dynamics of Multicellular Cell-Cycle Progression. Cell 132, 487–498.

Sánchez, I., and Dynlacht, B.D. (2016). Cilium assembly and disassembly. Nat. Cell Biol. 18, 711–717.

Satir, P., Pedersen, L.B., and Christensen, S.T. (2010). The primary cilium at a glance. J Cell Sci 123, 499–503.

Schmidt, R.L., Park, C.H., Ahmed, A.U., Gundelach, J.H., Reed, N.R., Cheng, S., Knudsen, B.E., and Tang, A.H. (2007). Inhibition of RAS-Mediated Transformation and Tumorigenesis by Targeting the Downstream E3 Ubiquitin Ligase Seven in Absentia Homologue. Cancer Res. 67, 11798–11810.

Seeley, E.S., Carrière, C., Goetze, T., Longnecker, D.S., and Korc, M. (2009). Pancreatic Cancer and Precursor Pancreatic Intraepithelial Neoplasia Lesions Are Devoid of Primary Cilia. Cancer Res. 69, 422–430.

Sfakianos, J., Togawa, A., Maday, S., Hull, M., Pypaert, M., Cantley, L., Toomre, D., and Mellman, I. (2007). Par3 functions in the biogenesis of the primary cilium in polarized epithelial cells. J. Cell Biol. 179, 1133–1140.

Simon, M.A. (1994). Signal Transduction during the Development of the Drosophila R7 Photoreceptor. Dev. Biol. 166, 431–442.

Singh, S., Howell, D., Trivedi, N., Kessler, K., Ong, T., Rosmaninho, P., Raposo, A.A., Robinson, G., Roussel, M.F., Castro, D.S., et al. (2016). Zeb1 controls neuron differentiation and germinal zone exit by a mesenchymal-epithelial-like transition. ELife 5, e12717.

Stebbins, J.L., Santelli, E., Feng, Y., De, S.K., Purves, A., Motamedchaboki, K., Wu, B., Ronai, Z.A., Liddington, R.C., and Pellecchia, M. (2013). Structure-Based Design of Covalent Siah Inhibitors. Chem. Biol. 20, 973–982.

Stecca, B., Mas, C., Clement, V., Zbinden, M., Correa, R., Piguet, V., Beermann, F., and Altaba, A.R. i (2007). Melanomas require HEDGEHOG-GLI signaling regulated by interactions between GLI1 and the RAS-MEK/AKT pathways. Proc. Natl. Acad. Sci. 104, 5895–5900.

Trivedi, N., Stabley, D.R., Cain, B., Howell, D., Laumonnerie, C., Ramahi, J.S., Temirov, J., Kerekes, R.A., Gordon-Weeks, P.R., and Solecki, D.J. (2017). Drebrin-mediated microtubule–actomyosin coupling steers cerebellar granule neuron nucleokinesis and migration pathway selection. Nat. Commun. 8, 14484.

Uziel, T., Zindy, F., Xie, S., Lee, Y., Forget, A., Magdaleno, S., Rehg, J.E., Calabrese, C., Solecki, D., Eberhart, C.G., et al. (2005). The tumor suppressors Ink4c and p53 collaborate independently with Patched to suppress medulloblastoma formation. Genes Dev. 19, 2656–2667.

Wechsler-Reya, R.J., and Scott, M.P. (1999). Control of Neuronal Precursor Proliferation in the Cerebellum by Sonic Hedgehog. Neuron 22, 103–114.

Xie, J., Murone, M., Luoh, S.-M., Ryan, A., Gu, Q., Zhang, C., Bonifas, J.M., Lam, C.-W., Hynes, M., Goddard, A., et al. (1998). Activating Smoothened mutations in sporadic basal-cell carcinoma. Nature 391, 90.

Yang, Z.J. (2008). Medulloblastoma can be initiated by deletion of Patched in lineage-restricted progenitors or stem cells. Cancer Cell 14, 135–145.

Yoon, J.W., Liu, C.Z., Yang, J.T., Swart, R., Iannaccone, P., and Walterhouse, D. (1998). GLI Activates Transcription through a Herpes Simplex Viral Protein 16-Like Activation Domain. J. Biol. Chem. 273, 3496–3501.

